# A miR-137-related biological pathway of risk for Schizophrenia is associated with human brain emotion processing

**DOI:** 10.1101/2020.08.03.230227

**Authors:** G. Pergola, A. Rampino, P. Di Carlo, A. Marakhovskaia, T. Quarto, L. Fazio, M. Papalino, S. Torretta, N. Amoroso, M. N. Castro, E. Domenici, J. Dukart, J. Khlghatyan, A. Monaco, T. Popolizio, R. Romano, L. Sportelli, H. Zunuer, G. Blasi, J.M. Beaulieu, A. Bertolino

## Abstract

Genome-Wide-Association studies have involved miR-137 in schizophrenia. However, the biology underlying this statistical evidence is unclear. Statistical polygenic risk for schizophrenia is associated with working memory, while other biological evidence involves miR-137 in emotion processing. We investigated the function of miR-137 target schizophrenia risk genes in humans.

We identified a prefrontal co-expression pathway of schizophrenia-associated miR-137 targets and validated the association with miR-137 expression in neuroblastoma cells. Alleles predicting greater co-expression of this pathway were associated with greater prefrontal activation during emotion processing in two independent cohorts of healthy volunteers (N_1_=222; N_2_=136). Statistical polygenic risk for schizophrenia was instead associated with prefrontal activation during working memory.

A co-expression pathway links miR-137 and its target genes to emotion processing and risk for schizophrenia. Low prefrontal miR-137 expression may be related with SCZ risk via increased expression of target risk genes, itself associated with increased prefrontal activation during emotion processing.

## Introduction

Inter-individual variation in traits with complex heritability, like psychiatric disorders, is associated with polygenic inheritance. As genes do not code for diseases, polygenic risk for psychiatric disorders like Schizophrenia (SCZ) and Autism Spectrum Disorders is likely enacted via molecular pathways affecting neurophysiological phenotypes (*1-3*). Accordingly, Polygenic Risk Scores cumulating the statistical effects of genetic variants associated with SCZ are associated with neural activity during working memory (*4-6*) and emotion processing (*7, 8*). This evidence indicates that statistical measures of genetic risk for SCZ are related with normal variation in brain features, including functional activity during cognitive and emotion processing as a possible mechanism linking risk to behavior. However, Polygenic Risk Scores do not provide information about the molecular pathways involved in the association of SCZ positive variants with system-level phenotypes. The purpose of whole-genome Polygenic Risk Scores lumping all variants together is to collect signal from all biological pathways involved. Understanding the mechanisms behind SCZ biology requires approaches integrating biological information in the score computation.

As many of the SCZ risk variants are non-coding and may control gene expression (*9-11*), co-expression regulation represents a potential mechanism to parse statistical summaries of genetic risk into biologically meaningful pathways. Importantly, several studies have found that SCZ risk genes are significantly co-expressed (*12-14*). PsychENCODE and other studies have suggested that biological networks gather converging effects of many genetic variants affecting neurophysiological and clinical phenotypes via expression quantitative trait loci (*11, 15-17*).

Candidate master regulators linking risk variants with gene expression include micro-RNAs targeting SCZ risk genes (*18*). Of these, miR-137 is probably the most biologically plausible candidate, potentially linking co-expression regulation with SCZ risk factors. Indeed, miR-137 genetic variation is associated with SCZ (*19*), with miR-137 brain transcription levels, with inter-individual variability in SCZ clinical measures, and with prefrontal cortex activity and connectivity in healthy controls (*20*). The risk allele of rs1625579, a miR-137 single nucleotide polymorphism (SNP) associated with SCZ, is associated with lower miR-137 expression in the dorsolateral prefrontal cortex (*21*), with working memory-related prefrontal activity (*22, 23*) and with activity and connectivity of the emotion processing brain network (*24, 25*). Taken together, these studies suggest that miR-137-related risk for SCZ is relevant to brain function during working memory and emotion processing, possibly via pleiotropic effects on SCZ phenotypes.

MiR-137 regulates the expression of many genes (*26*), therefore one possible mechanism for its effects in SCZ is co-expression regulation. Accordingly, two studies parsed the polygenic effects of miR-137 on task-based brain activation.

Potkin and coworkers (*27*) reported that variants associated with brain activity during WM are located in loci co-expressed with SCZ risk genes. Consistent with the role of miR-137 in orchestrating working memory gene co-expression, miR-137 target genes harbored variants associated with brain activity during working memory. However, there was no direct link between miR-137 targets and risk-related gene co-expression in the human brain.

Cosgrove and coworkers (*28*) parsed the SCZ-Polygenic Risk Score by including only miR-137 targets and found that it was associated with brain activity during working memory. They failed to identify significant effects of the Polygenic Risk Score in 83 participants performing an emotion processing task. While these studies are in agreement on the relationship between miR-137 targets, SCZ risk, prefrontal cortex, and working memory, they do not support a functional role of miR-137 in emotion processing. This evidence suggests that miR-137 is related with SCZ via pathways common to working memory, but it is worth reiterating that Polygenic Risk Scores do not provide biological information. The question thus stands, whether miR-137 is linked with working memory *per se*, or whether these findings instead depend on weighting genetic variants by SCZ risk.

At variance with the evidence derived from these approaches, molecular evidence independent of SCZ implicates miR-137 in emotion processing. MiR-137 is expressed in the amygdala more than in the prefrontal cortex (*21*), and its expression is upregulated by the metabotropic glutamate-receptor-5 (mGluR5); in turn, miR-137 contributes to mGluR5 function in the emotion processing brain network (*20, 29*). The lacking association between miR-137 and emotion processing in studies employing polygenic approaches may be attributed to statistical power limitations of earlier studies. Alternatively, the polygenic approaches used may have diluted the effect of risk pathways affecting emotion processing with unrelated signal, e.g., because ***only a subset of the SCZ loci targeted by miR-137 is associated with emotion processing***. We thus hypothesized that 1) miR-137 target genes are co-expressed in the human brain with SCZ risk genes, and 2) are associated with functional brain activity during emotion processing.

Therefore, we investigated whether: **i)** miR-137 target genes are co-expressed in the human prefrontal cortex and how they relate with SCZ risk; **ii)** co-expressed miR-137 targets are related with working memory or emotion processing brain activity in humans based on biologically stratified genetic scores, and ***which genes*** are involved; **iii)** SCZ Polygenic Risk Scores combining variants harbored in miR-137 target genes based on SCZ risk stratification predict working memory or emotion processing brain activity in larger cohorts than those previously reported. Figure 1 reports a synopsis of the study.

**Figure 1.**
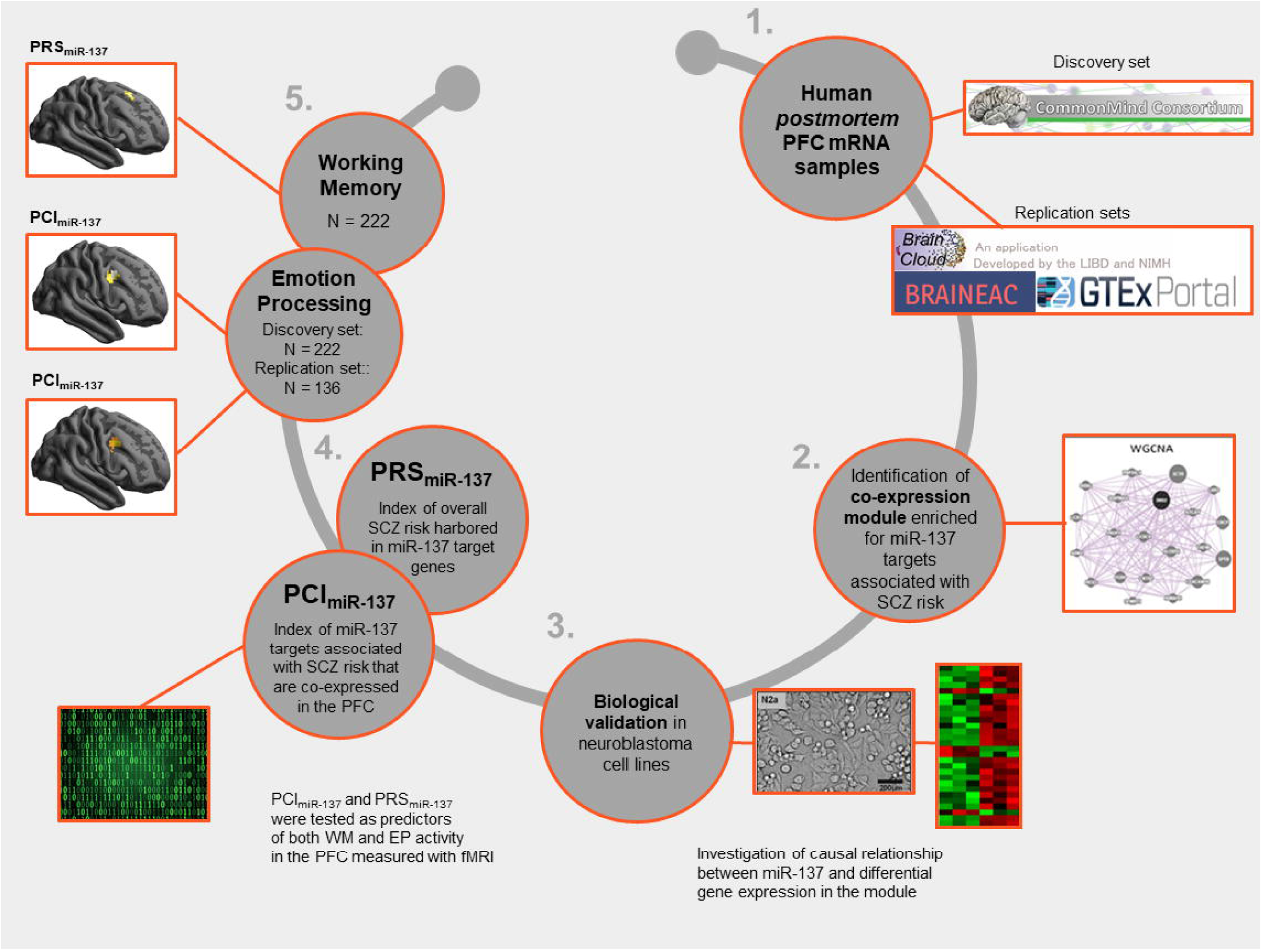
Study design.

We identified co-expression modules enriched for miR-137 targets in human *post-mortem* prefrontal cortex and validated the modules in neuroblastoma cells. We linked *post-mortem* co-expression networks with system-level phenotypes via co-expression quantitative trait loci (co-eQTLs (*30-32*)), i.e., SNPs associated with gene co-expression which were combined into a Polygenic Co-expression Index (PCI_miR-137_) used to test further associations. The PCI_miR-137_ indexes *the predicted expression of a subset of miR-137 targets associated with SCZ risk that are co-expressed in the prefrontal cortex*. Therefore, this index identifies a miR-137 related biological pathway of risk.

Additionally, we computed a previously reported Polygenic Risk Score parsed to include variants harbored in miR-137 target genes (*28*). The PRS_miR-137_ includes the *overall statistical risk for SCZ harbored in miR-137 target genes*, weighted by their GWAS-derived effect size. Therefore, it is an index of miR-137 related statistical measure of genetic risk. Both PCI_miR-137_ and PRS_miR-137_ were tested as predictors of both working memory and emotion processing activation in the prefrontal cortex assessed via fMRI.

## Results

Our gene co-expression network included 51 modules, plus the *grey* module of 3,018 non-clustered genes. Permutation tests revealed that gene-gene relationships in 28 out of 51 modules were preserved in three replication datasets (p<α_Bonferroni_ = .00033; Figure S1). Gene loadings on the module expression were concordant across all replication datasets (binomial p-values, all p<10^−12^; Figure S2), suggesting that the replications identified gene-gene relationships in the same direction as the discovery set. Four of these 28 modules were enriched for miR-137 target genes (p<α_Bonferroni_=0.05/(28×4)=.00045; Table S3). Out of these four, only *Darkorange* was significantly enriched for genes in the SCZ risk loci (p=.0052 with α_Bonferroni_=0.05/4=0.0125; nine genes in nine risk loci, see Table 1; Table S4 reports the list of *Darkorange* genes). Using a different method to quantify SCZ association, we found that genetic variants harbored within *Darkorange* genes were associated with greater SCZ risk compared to all remaining modules (MAGMA (*33*); p=.0012 with α_Bonferroni_=0.05/28=0.0018; Figure 2). The module eigengene, i.e., the first principal component of *Darkorange* gene expression, explained the majority (54%) of *Darkorange* expression variance. Overall, the over-representation of genes and variants in the risk loci suggested that a component of the genetic risk for SCZ converged into *Darkorange*.

**Table 1.**
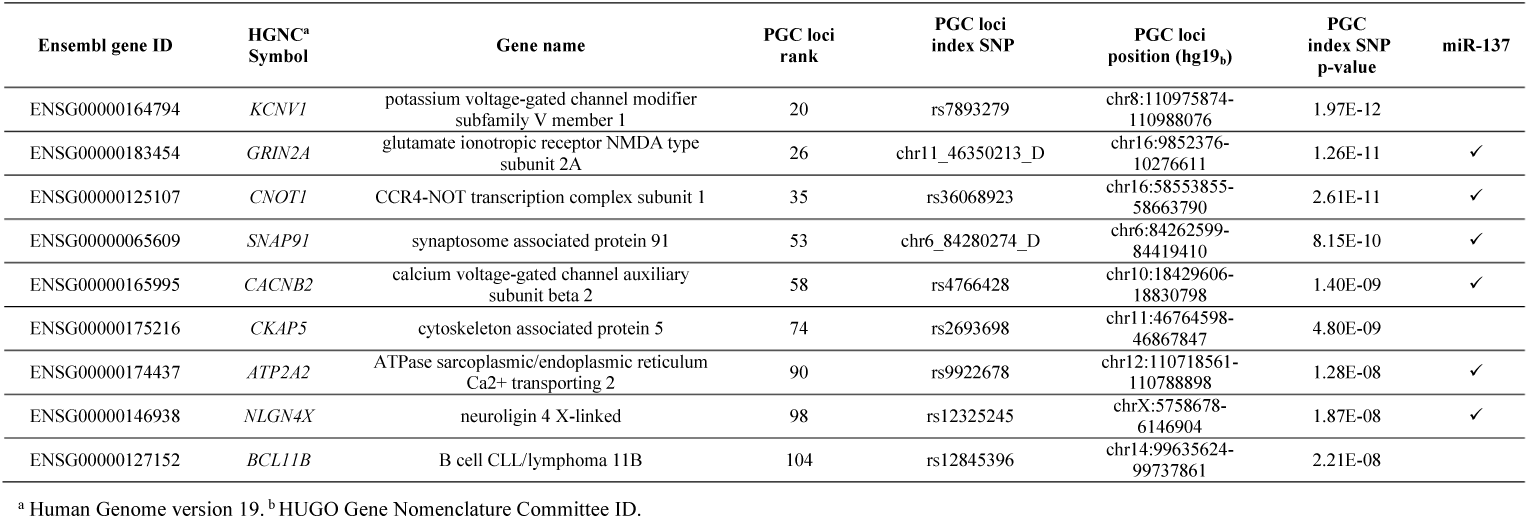
PGC loci and genes overlapping with the module Darkorange and miR-137 targets.

**Figure 2.**
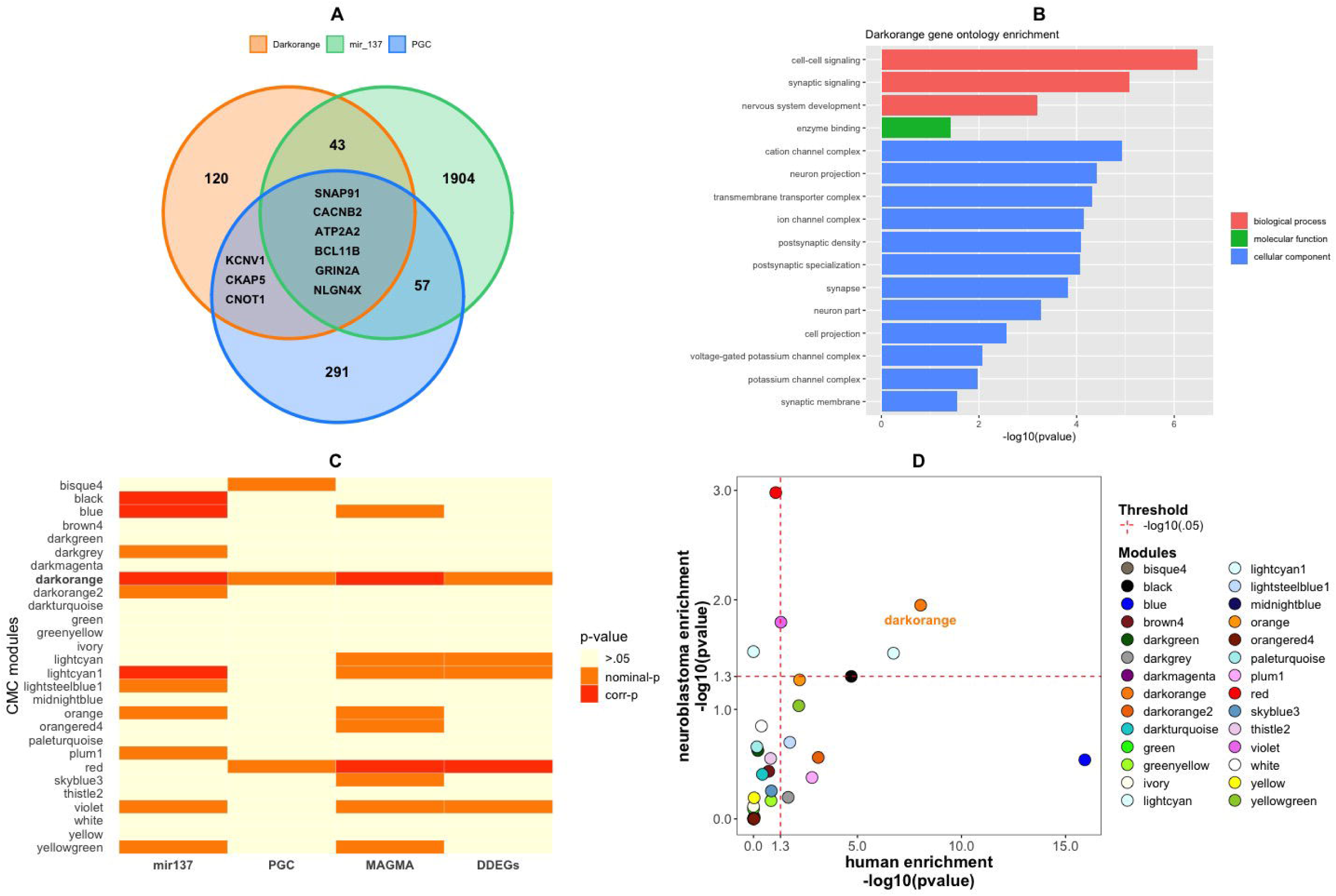
Module prioritization. Features of the *Darkorange* module. **A**. Intersection between *Darkorange* genes, Psychiatric Genomic Consortium (PGC) SCZ genes, and miR-137 targetome bioinformatic predictions in humans. **B**. Gene ontologies associated with *Darkorange* genes. Reported p-values are corrected for multiple comparisons. **C**. Overview of the over-representation in all replicated modules (28 out of 51) of miR-137 predicted targets in humans, of SCZ genes, of genetic variants associated with SCZ (MAGMA), and of dose-dependent expressed genes in neuroblastoma cells (DDEGs). The colormap highlights the specificity of *Darkorange* relative to other modules. **D**. Scatter plot illustrates the enrichment of each replicated module for miR-137 targets as assessed in neuroblastoma cells and in bioinformatic predictions in humans. The two assessments co-vary except for outliers *Blue* and *Red*.

### Darkorange is enriched for neuronal markers and gene ontologies relevant to schizophrenia

*Darkorange* was comprised of 173 genes (156 protein-coding). *Darkorange* presented marker gene expression characteristic of glutamatergic and GABAergic neurons (*34*) (Figure S3) and was functionally enriched for biological processes relevant to SCZ (Figure 2), including synaptic signaling (GO:0099536, 19 genes, fold enrichment=5.8, p_corrected_=8.37×10^−6^) and nervous system development (GO:0007399, 41 genes, fold enrichment=2.4, p_corrected_=6.39×10^−4^). No chromosomal locus was significantly over-represented in *Darkorange*. Specific Expression Analysis revealed that *Darkorange* genes are preferentially expressed in the PFC during young adulthood (p_corrected_=1.7×10^−5^, [pSI<.01]), with a pattern closely matching published findings on SCZ genes (*35*) (Figure S4). *Darkorange* was not enriched for TWAS (all p>.05).

### Darkorange topology links schizophrenia risk genes with miR-137 targets

Brainspan data confirmed that miR-137 expression was more correlated with miR-137 targets than non-targets (Wilcoxon rank-sum test, p = .044). Specifically, correlations were more negative for miR-137 targets (on average, r = -.61) than for non-targets (average r = -.47). These results indicate that the genes identified above as being co-expressed with miR-137 are indeed more associated than others with miR-137 expression.

Within *Darkorange*, the analysis of intramodular connectivity showed that miR-137 targets were more central, i.e., more strongly connected with the rest of the module, than non-targets (Wilcoxon rank-sum tests, p = 0.0093). Eight out of nine SCZ risk genes (Figure 2) had higher correlations with miR-137 targets than non-targets (all p < .05); gene *KCNV1* reached marginal significance (p = .066). These results indicate that miR-137 targets are connected to SCZ risk genes above chance level; SCZ risk genes are more related with targets than with non-targets within *Darkorange*.

### Darkorange is enriched for genes modulated by miR-137

To assess directly the contribution of miR-137 to the regulation of *Darkorange*, we used a cellular system allowing for a titration of miR-137 expression. Plasmid transfections were used to overexpress (OE) miR-137 in a Neuro2A (N2A) neuroblastoma cells. A CRISPR/Cas9 approach (Figure S5) was also developed to disrupt (KO) miR-137 expression in these same cells. Evaluation of miR-137 expression using quantitative PCR confirmed the efficacy of the KO as well as OE procedures (Table S5). We quantified the effect of various levels of miR-137 expression on genome-wide gene expression using microarray analysis (Supplemental Information [SI]-1.3).

We identified 580 dose-dependently expressed genes (DDEGs) showing linear covariation with miR-137 across the transcriptome (q_FDR_<.05). *Darkorange* included 13 DDEGs out of 141 genes expressed in neuroblastoma cells, a significant proportion (p=.011; Figure 2; Figure S6). We assessed whether any other replicated modules were significantly enriched as a negative control. Only few modules over-represented miR-137 DDEGs (Figure 2), consistent with the human *post-mortem* data and supporting the relative specificity of *Darkorange* for miR-137 targets.

### The Polygenic Co-expression Index combines reproducible associations of SNPs with Darkorange

We translated *Darkorange* gene co-expression into the PCI_miR-137_ via the association of genetic variants with the module eigengene. SNP weights are available in Table S6. PCIs including between 7 and 15 SNPs were replicated (BRAINEAC: one-tailed p<.05, Table S6). Table 2 includes annotations of the first 15 SNPs. We adopted an inclusive approach to collect signal from multiple module genes and therefore tested the PCI_miR-137_ including 15 SNPs in the neuroimaging sample.

**Table 2.**
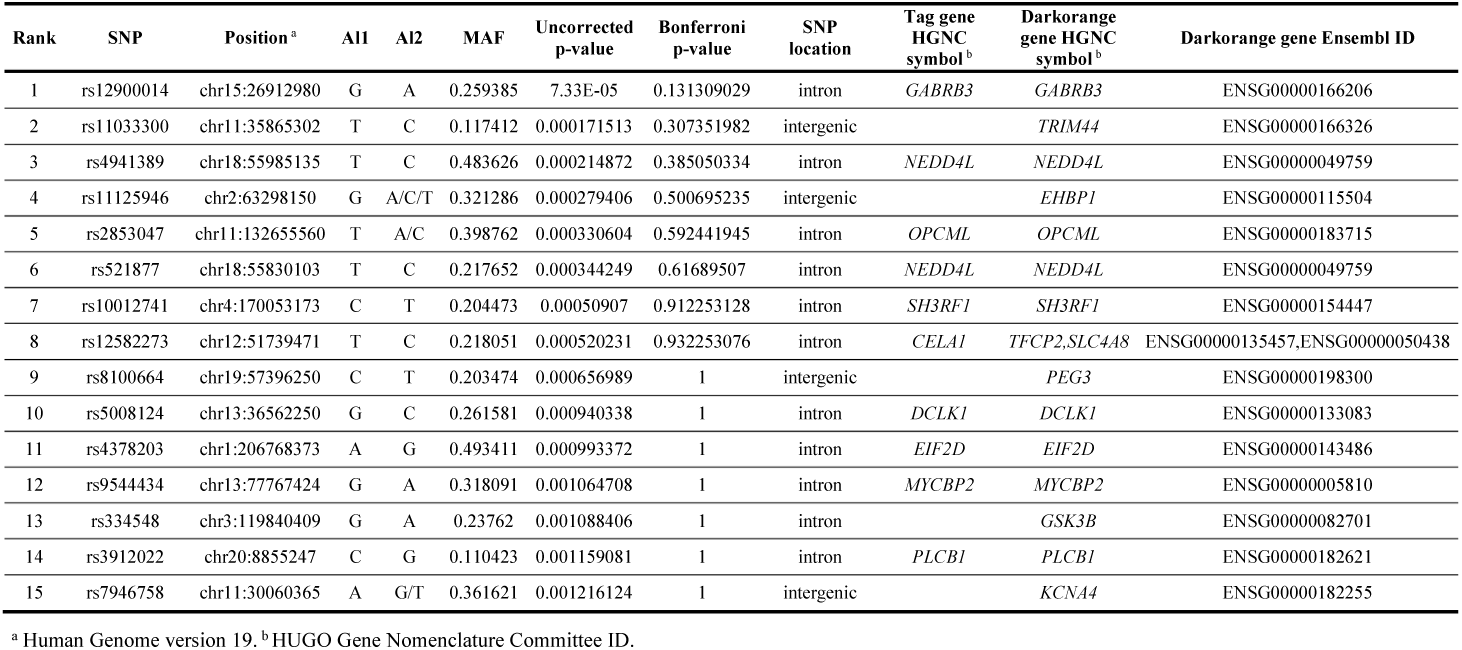
SNPs associated with the module *Darkorange*.

### Neuroimaging results confirm the association of polygenic risk for schizophrenia with working memory but reveal an association of miR-137 related co-expression with emotion processing

Table S7 reports complete fMRI statistics. During the emotion processing task, the linear term of the PCI_miR-137_ correlated positively with BOLD activity in a right prefrontal cluster including Brodmann areas 8, 9, and 46 within the discovery sample (peak Z=5.1; p_FWE_=.001; 97 voxels; MNI coordinates x=50; y=12; z=42; Figure 3). In the replication sample, the effect was in the same direction and significant in BA9 (peak Z=3.5; p_SVC_=.007; 42 voxels; MNI coordinates x=46; y=12; z=31). Therefore, greater *Darkorange* predicted co-expression was associated with greater prefrontal activity during emotion processing. Greater PCI_miR-137_ was also associated with lower prefrontal-amygdala connectivity during emotion processing (p_SVC_ < .05; Figure S7). We found no effect of PRS_miR-137_ on emotion processing.

**Figure 3.**
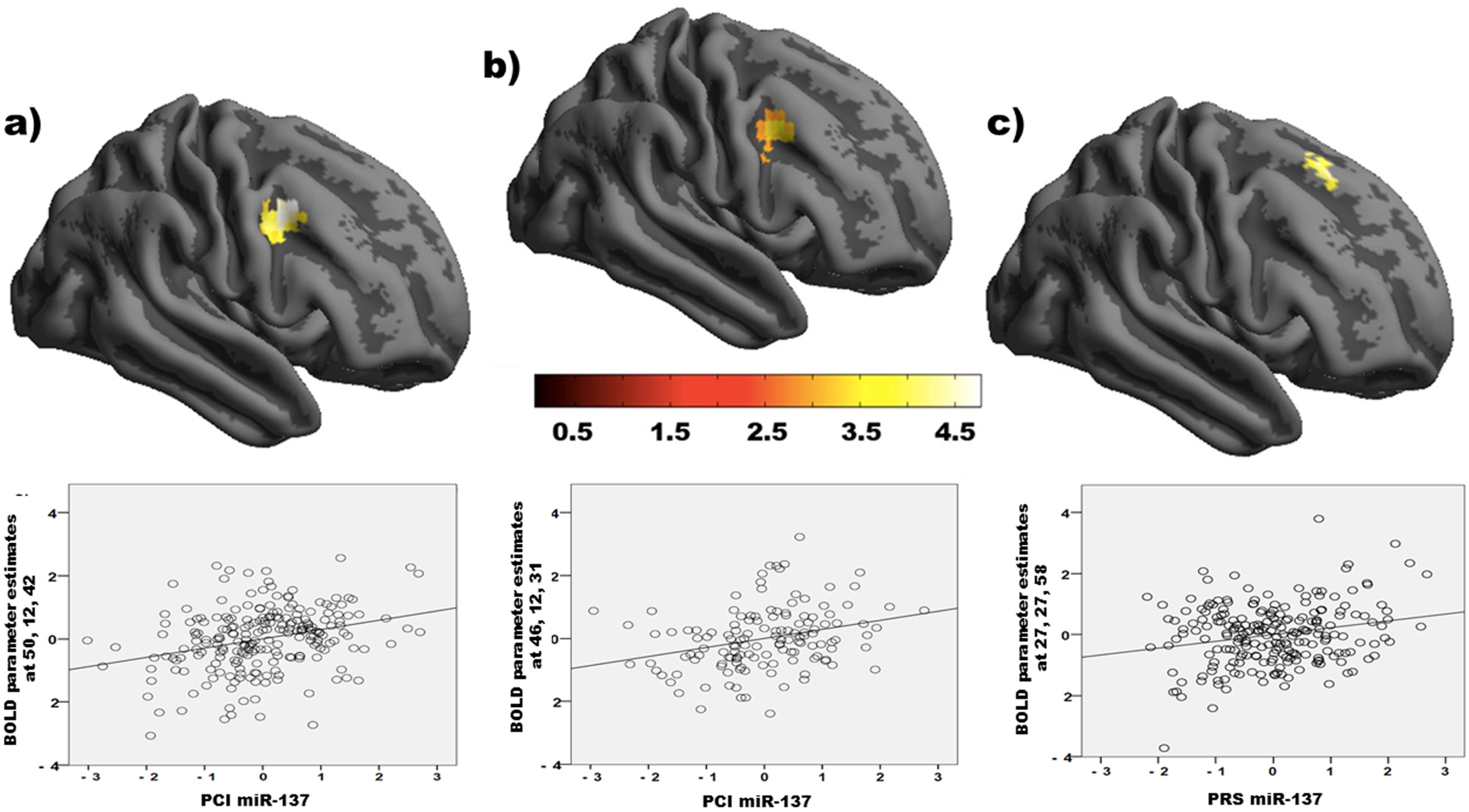
fMRI results. Top panels: render of the brain activation associated with PCI_miR-137_ during EP in the discovery (**A**.) and replication sample (**B**.). **C**. Render of the brain activation associated with PRS_miR-137_ during working memory in the discovery sample. Color bar indicates Z scores. Right in the figures is right in the brain. Each scatterplot refers to the cluster highlighted in the render on its top. In the scatterplots, axes are scaled with mean = 0 and standard deviation = 1.

Conversely, we found no significant effect of the PCI_miR-137_ on working memory-related activity. Instead, PRS_miR-137_ computed with two of the three SCZ association thresholds (p<.05 and p<.5) correlated positively with working memory activity in a right prefrontal cluster including BA8 (SNP association p<.05 threshold: peak Z=4.5; p_FWE_=.008; 37 voxels; MNI coordinates x=27; y=27; z=58). This effect was specific for PRS_miR-137_ and was not observed when using SCZ risk variants not harbored within miR-137 targets (SI-2). Further analyses suggested that statistical power was not an issue for the PCI_miR-137_ n-back analysis and removing *Darkorange* genes from the PRS_miR-137_ did not affect results (SI-2). Additionally, to establish the specificity of the imaging associations, we explored ththe sameis association with PCIs derived from other modules enriched for SCZ risk genes (*Black, Blue*, and *Lightcyan*; see Figure 2) as a negative control. To compute these scores, we used the top 15 independent SNPs within the module genes like we did for the *Darkorange* PCI. We found no significant association with the n-back task and no replicated association with the Faces task (data available on request).

## Discussion

We hypothesized that miR-137 regulates the co-expression of SCZ risk genes in the human prefrontal cortex and evaluated the involvement of miR-137 targets in working memory and emotion processing. Indeed, we found that miR-137 targets are co-expressed in at least four reproducible co-expression modules. Of these, only *Darkorange* shows over-representation of SCZ genes and has a link with miR-137 supported by *in vitro* evidence. *SNAP91, CACNB2, ATP2A2, BCL11B, GRIN2A*, and *NLGN4X* genes appear at the intersection between miR-137-associated co-expression and SCZ risk. Gene Ontology analysis links *Darkorange* genes with Neurodevelopment and Synaptic Signaling, two biological functions implicated in SCZ. When we translated miR-137 effects on its targets into two polygenic indices respectively reflecting co-expression of a subset of SCZ risk genes targeted by miR-137 (PCI_miR-137_) or the whole-genome association with SCZ risk (PRS_miR-137_), we obtained differential neuroimaging readouts in neurotypical individuals. We replicated prior evidence that PRS_miR-137_ is associated with prefrontal activity during working memory, confirming that SCZ-weighted genetic score associate miR-137 with working memory. Instead, genetic scores based on co-expression tell a different story: greater ***PCI***_***miR-137***_ ***derived from a subset of miR-137 targets co-expressed in the human PFC*** is associated with greater prefrontal activity and lower prefrontal-amygdala coupling during emotion processing, but is not related with working memory. None of the three modules enriched for miR-137 but not enriched for SCZ showed an association with working memory. Taken together, this evidence shows that miR-137 targets are involved both in working memory and emotion processing SCZ phenotypes, but a subset of miR-137 potentially co-regulated genes appears more closely associated with emotion processing.

Our approach has two advantages: first, it incorporates SCZ GWAS information in the module prioritization, but also includes co-expression information – whereas the Polygenic Risk Score only includes GWAS information; second, it parses miR-137 targets into sets of co-expressed genes, whereas the Polygenic Risk Score computed with all miR-137 targets cumulates statistical effects with parsing them. Dissecting the overall genetic risk for SCZ into biologically meaningful information related with miR-137 co-expression regulation reveals previously unreported links between miR-137 targets and *in vivo* phenotypes of SCZ. We suggest that the functional role of miR-137 in emotion processing is mediated by the co-expression of a gene set involved in SCZ risk.

### Common variation in target genes of miR-137 is related with emotion processing

We show that miR-137 is associated with co-expression of SCZ risk genes in human non-psychiatric *post-mortem* PFC. Results are consistent across independent samples, thus reducing the chance of type I error (*3, 36*). *In silico* predictions of miR-137 association with *Darkorange* co-expression are supported by *in vitro* experiments. Prior evidence that miR-137 SCZ-associated variants are related with working memory and not emotion processing brain activity is replicated, and we identify a role of miR-137 in emotion processing. Importantly, the sizes of the fMRI cohorts previously tested (*25, 28*) ranged up to 86 participants, whereas here we studied 358 individuals overall. As we assessed healthy individuals without familial psychiatric history, our findings speak to the role of physiological variation in a subset of miR-137 targets. In particular, greater miR-137 target gene expression (mostly associated with lower miR-137 expression in the Brainspan dataset and in our experimental validation – Figure S6) – in turn linked with greater SCZ risk (*21*) – is associated with greater prefrontal activity and lower prefrontal-amygdala coupling during emotion processing. These findings are consistent with prior evidence of greater prefrontal activity in individuals carrying rs1625579 risk allele (*23*) and with decreased prefrontal-amygdala connectivity associated with SCZ risk (*37*). Notably, our findings link lateral prefrontal cortex with amygdala, rather than ventromedial prefrontal regions investigated in previous studies (*25*), like in previous reports associating disrupted lateral prefrontal-amygdala connectivity with genetic risk for SCZ (*38*). We propose that lower miR-137 expression in the prefrontal cortex is associated with risk for SCZ by an increase of target gene expression. Some of the SCZ risk genes are co-expressed in the pathway we identified here, which is reproducibly related with increased prefrontal activity during emotion processing and decreased prefrontal-amygdala coupling.

### Co-expression of miR-137 target genes

The over-representation of synaptic and neurodevelopmental genes in *Darkorange* is consistent with prior evidence about miR-137 functions (*39, 40*). For example, He and coworkers (*41*) found that miR-137 expression was associated with synaptic protein levels in mouse hippocampus relating such dysfunction with Synaptotagmin-1, the main calcium sensor regulating brain synaptic vesicle release (*42*). In mice, overexpression of the Synaptotagmin-1 coding gene *Syt1* antagonizes deficits caused by miR-137 OE (*43*). Therefore, it is noteworthy that *Syt1* is a member of *Darkorange* and a DDEG.

Two further relevant genes are *GRM5* and *GSK3B. GRM5*, coding for mGluR5, has been implicated in SCZ (*44*) and in affective phenotypes (*45*). Interestingly, mGlu5 acts via a pathway mediated by activation of phospholipase C and protein kinase C (*46*), a pathway also represented in *Darkorange* (*PLCB1* and *PRKCB*). Similarly, many studies have implicated the Glycogene-Synthetase-Kinase-3beta (*GSK3B*) gene in emotion processing (*47*) and in SCZ (*48*). *GSK3B* ranks among the most connected genes within *Darkorange* (Table S4), making it a potential hub of co-expression. These findings are consistent with prior reports implicating *GSK3B* variants in SCZ intermediate phenotypes (*48-50*).

### Limitations

We identified *Darkorange* based on over-representation of predicted miR-137 targets. About 50 targets have been experimentally validated (*26*), but the number is likely much larger. As parsing 50 genes across 51 co-expression modules would be inadequate for enrichment analysis, we relied on predictions, as done elsewhere (*28*). The reliability of these predictions limits our inferences. We endeavored to address this limitation by *in vitro* experiments showing that *Darkorange* stands out at the intersection between bioinformatics and findings in neuroblastoma cells (Figure 2). Notably, the overexpression we employed may be above the physiological range of variation, hence potentially exceeding the physiological response to miR-137. Although we do not have evidence that DDEGs are directly, rather than indirectly, targeted by miR-137, this evidence suggests a miR-137 association with *Darkorange*. Finally, relative to our previous work (*31*), we traded sample size for homogeneity, by selecting only neurotypical individuals of Caucasian ancestry in order to match the SCZ-GWAS (*19*) and our fMRI cohort.

### Conclusions

As in prior reports, polygenic SCZ risk enriched for miR-137 target genes is associated with WM-related PFC activity. Furthermore, we identified the regulation of *Darkorange* gene co-expression as a mechanism of miR-137 involvement in risk for SCZ. *Darkorange* co-eQTLs were associated with prefrontal activity and connectivity during emotion processing, in line with previous knowledge about miR-137 and emotion processing. We propose that low miR-137 expression in the prefrontal cortex is associated with SCZ risk via the increased expression of specific risk genes which is associated with increased prefrontal activation during emotion processing.

The study of co-expression in *post-mortem* tissue and neuroimaging phenotypes disentangles the functional genetics of gene regulators with clinical relevance beyond PRS approaches. The biological role of miR-137 in SCZ likely acts via co-expressed gene sets. Here we have identified one of such sets related with emotion processing brain activity and connectivity in humans.

## Materials and Methods Experimental Design

### Identification of a Gene Co-expression Network in the human prefrontal cortex

We used CommonMind Consortium RNA-sequencing data from *post-mortem* prefrontal cortex (*9*) to identify a gene co-expression network in HCs by means of Weighted Gene Co-expression Network Analysis (WGCNA)(*51*). A dataset comparable with the fMRI sample in terms of age and ancestry included 147 neurotypical Caucasian adults (demographics in Table S1; SI-1.1). After correcting the expression data for confounding variables, we computed WGCNA as previously reported (*31*) on 17,173 genes. Here we used Spearman’s correlation as a weighted measure of gene-gene relationships (*52*) to limit the impact of deviations from normality in expression data (*53*) (SI-1.1).

We validated the discovered gene-gene relationships using three independent datasets (BrainEAC (*54*) Frontal Cortex [N=123], Braincloud PFC BA46 [N=59](*13*) and GTEx (*55*) PFC BA9, [N=84], Table S1). A permutation approach served to compare module cohesiveness (*56*)(SI-1.1). We assessed the replication of each module in the three datasets and performed a meta-analysis to obtain module-wise replication p-values (sum-log Fisher’s method; α_Bonferroni_=.05/(N_modules_×3)).

### Module prioritization and functional analysis

We assessed the overlap between genes in the successfully replicated co-expression modules and miR-137 target genes derived via a meta-analysis of four miRNA target databases (*57-60*). Then, we assessed the over-representation of genes located in loci previously associated with SCZ at genome-wide significance (*19*) within the miR-137-enriched modules (Table S2; SI-1.2). We characterized the modules of interest using the Gene Ontology Database (PANTHER)(*61*). We investigated module function via enrichments for chromosomal locus (*62*), cell-type, and brain region-specific expression pattern (*63*) during neurodevelopment. To further assess the potential relationship between risk and gene expression, we also tested the over-representation of transcriptome-wide association study (TWAS)(*11, 17*, 64) variants in the selected modules.

### miR-137 targets in the prioritized modules and topology of schizophrenia risk genes

As there was no direct quantification of miR-137 in the dataset we used to compute the network, we used the Brainspan dorsolateral prefrontal cortex data to verify the association between miR-137 and the modules of interest (only N = 10 had a miR-137 quantification available). We computed Pearson’s correlations between miR-137 and each module gene; then, we assessed the correlation difference between miR-137 targets and non-targets to support the set of targets we used.

The fact that the prioritized modules show significant enrichment for both miR-137 and SCZ gene lists does not necessarily imply that the involved miR-137 targets are linked to SCZ genetic risk. Therefore, we overlapped the gene lists to identify miR-137 targets associated with SCZ risk within modules of interest and studied the topology of the network. Specifically, we explored whether miR-137 targets were connected to SCZ risk genes above chance level and whether SCZ risk genes were more related with targets than with non-targets in the prioritized modules. To this aim, we used intramodular connectivity, a standard output of WGCNA, and assessed the difference of connectivity between targets/non-targets and SCZ risk genes (Wilcoxon rank-sum tests). To test the relationship of SCZ risk genes with targets and non-targets, we computed module-wise correlation matrices and assessed the difference in correlations with Wilcoxon rank-sum tests.

### Experimental validation in neuroblastoma cells and data analysis

We modulated the expression of miR-137 in neuroblastoma Neuro2A cells to assess WGCNA-based predictions of a link between miR-137 and the expression of genes belonging to the prioritized modules (SI-1.3). A CRISPR-Cas9 approach involving a plasmid expressing both Cas9 and a specific small guide RNA was used to generate miR-137 KO cells. A PcDNA3.2/V5 mmu-mir-137 plasmid expressing miR137 was used for OE conditions (*65*). Impact of dose-dependent linear changes in miR-137 expression on transcriptome-wide expression was evaluated using microarrays (Mouse Gene 2.0 ST microarrays, Affymetrix/Thermo-Fisher). In the linear models, gene expression was the dependent variable and miR-137 quantification was the predictor. We thresholded results ad q_FDR_<.05 to obtain a list of dose-dependently expressed genes (DDEGs) and assessed DDEGs over-representation in the target modules (SI-1.3).

### Biological genetic stratification of miR-137 target genes: the Polygenic Co-expression Index

In order to test systems-level phenotypes associated with the module of interest, we generated a Polygenic Co-expression Index (PCI_miR-137_). We identified SNPs in the module genes associated with the first principal component of module gene expression (a measure of co-expression of the whole module) and combined them into the PCI_miR-137_(*13, 30, 66*)(SI-1.4). We estimated the effect of allelic dosage via a Robust Linear Model (*rlm* function - *robust* R package) and ranked SNPs according to their p-value (*31*). Following prior reports, we assessed the top 50 ranked SNPs for PCI computation (*31*), and computed the PCI_miR-137_ picking the top set of SNPs that replicated the association with co-expression in the largest replication dataset we had available (BrainEAC; SI-1.4).

### Neuroimaging study

#### Participants

We recruited 358 healthy volunteers distributed in a discovery cohort of 222 participants who had both EP and WM data available and a replication cohort of 136 participants for the emotion processing task (Table S1). We evaluated a single working memory cohort because the PRS_miR-137_ effect has already been reported independently (*28*) and the replication cohort did not have working memory scans. Inclusion/exclusion criteria have been described elsewhere (SI-1.5)(*67*). Participants signed an informed consent complying with the Declaration of Helsinki after full explanation of all procedures approved by the local ethics committee.

#### Genetic score computation

We genotyped all participants (*68*) and performed imputation using Sanger Imputation Service and 1000 Genomes Project Phase 3 reference panel (including pre-phasing and imputation with SHAPEIT2+PBWT (*69, 70*); genomic coordinates on GRCh37/hg19 genomic reference build). We filtered out SNPs with missing rate >.05, Hardy-Weinberg equilibrium p<10^−6^, and minor allele frequency (MAF)<.01 and computed genomic eigenvariates to control for population stratification (*71*).

The PCI_miR-137_ is the average of the co-expression effects of the top-ranked SNP alleles selected in the *post-mortem* study, which replicated in BrainEAC. Instead, we computed the PRS_miR-137_ for SCZ using standard procedures (*71*) with the miR-137 target gene list and SNPs reported in the original article (*28*)(SNP-level significance thresholds: p<10^−5^, p<.05, p<.5). SI-1.6 reports further scores computed to control for possible confounders and the pertinent results.

#### Neuropsychological tasks

Our emotion processing Faces task (*72-74*) presented participants with angry, fearful, happy and neutral facial expressions from a validated set of facial pictures (SI-1.5). The N-back task probes working memory and has been widely used in neuroimaging (*75-77*). Participants performed three runs of a block design version of the task: 1-Back vs. 0-Back; 2-Back vs. 0-Back, and 3-Back vs. 0-Back, each lasting 240 s.

### Statistical Analysis

fMRI data collection, pre-processing, and analysis followed standard procedures (SI-1.5)(*13*). We used SPM12 to perform multiple regression analyses for both tasks and for the two samples separately. For the co-expression analysis, we used the linear and quadratic terms of PCI_miR-137_ (*30, 66, 78*) as predictors and age, gender, and five genomic eigenvariates as covariates. For the PRS_miR-137_ analyses, we used PRSs as predictors and the same covariates reported above. In the working memory/emotion processing sample, we report results surviving p_FWE_<.05 threshold at whole brain level masked by task activity. In the emotion processing replication sample, we used the cluster detected in the discovery analysis and performed a small volume correction (p_SVC_<.05).

To link miR-137 target gene predicted expression in the prefrontal cortex with prefrontal-amygdala functional coupling, we additionally performed a connectivity analysis using a genetic-physiological interaction approach (*79*). Specifically, we used the prefrontal seed associated with the PCI_miR-137_ and extracted movement-and task-corrected estimates from the bilateral amygdala (WFU Pickatlas (*25*); SI 1.5).

## Acknowledgments

CommonMind Consortium data were generously provided to GP by the NIMH and CommonMind Consortium. Data acquisition was possible thanks to the work by Dr. Linda A. Antonucci, Dr. Barbara Gelao, Dr. Marina Mancini, Dr. Annamaria Porcelli, and Dr. Paolo Taurisano. We gratefully acknowledge the work by Prof. Roberto Bellotti and Dr. Sabina Tangaro (Department of Physics – University of Bari Aldo Moro), Dr. Lucia Colagiorgio, Dr. Anna Monda, Annalisa Lella and Elisabetta Volpe (Department of Basic Medical Science, Neuroscience, and Sense Organs – University of Bari Aldo Moro), who contributed to data analysis.

