## Supplementary Material for "A miR-137-related biological pathway of risk for Schizophrenia is associated with human brain emotion processing"

***Supplemental Information (SI)***

**This file includes:**

**1. SI Materials and Methods**

1.1. Gene Co-expression Network
1.2. Module Prioritization
1.3. Experimental Validation and Data Analysis
1.4. Polygenic Co-expression Index
1.5. Neuroimaging Study
1.6. Control imaging genetics analyses

**2. SI Results**

**3. SI Figures**

Figure S1. Module-wise replication

Figure S2. Gene loading replication on *Darkorange* module eigengene

Figure S3. Module-wise cell specificity

Figure S4. Brain region specific expression analysis

Figure S5. TIDE analyses of CRISPR-CAS9 editing efficiency for g1 and g3 in Neuro2A cells.

Figure S6. Darkorange Dose-dependent expressed genes across experimental conditions.

Figure S7. Association of the Polygenic Co-expression Index with prefrontal-amygdala connectivity.

**4. SI Tables**

Table S1. Demographics

Table S2. Genes in the PGC list included in the network

Table S3. The CMC network: enrichment statistics for schizophrenia loci and mir137 targets

Table S4. *Darkorange* genes and overlap with PGC, mir137 targets and mir37 DDEGs

Table S5. Experimental validation design

Table S6. SNP weights

Table S7. fMRI statistics

**5. SI References**

1. **SI Materials and Methods**

**1.1 Gene Co-expression Network**

In order to reduce the differences between the postmortem and the fMRI datasets, in the CommonMind Consortium (CMC) dataset and in all replication samples we selected adult subjects (> 17 years) of Caucasian ancestry (as determined per the CMC meta-data (1)); additionally, RNA Integrity Number (RIN) ≥ 7.0 guaranteed the use of high-quality observations. After applying these filters, 163 out of the initial 279 HCs were included in our sample. Sample sizes were very unbalanced across collection sites (MSSM = 74, PITT = 73, PENN = 16), thus we removed the smallest sample (PENN) to reduce site-to-site heterogeneity.

Gene-level RNA expression was quantified as Reads Per Kilobase per Million mapped reads (RPKMs); see for genotyping, reference, and genetic procedure details (1). We selected well expressed genes above median RPKM ≥ 3 and log-transformed RPKM values with an offset of 1, i.e., log_2_(RPKM+1). Since RNA expression data are affected by systematic noise (2), we preprocessed the expression matrix correcting the expression of each gene for explicit confounders, i.e., the linear and quadratic effects of RIN, post mortem brain tissue pH, age, post mortem interval, institute, gender, and library batch as computed in the original CMC publication (1) by means of general linear models. We then computed an unsigned co-expression network to keep the information regarding negative correlations between genes that may be meaningful for regulatory genetic elements (3). The correlation matrix was further transformed into an adjacency matrix by raising Spearman’s coefficients to a positive exponent, β = 7, selected to ensure the identification of a “scale-invariant” network *(4,* 5). Hierarchical clustering then grouped genes into sets called “modules” (arbitrarily labeled with colors), with the “grey” module representing non-clustered genes. The module eigengene (*ME*), i.e., the first principal component of the expression of genes within each module (i.e., co-expressed gene set), indexed gene expression covariation *(6-*10).

We sought to replicate network analysis results by using an approach that tests whether the topological relationships between genes in the second dataset mirror those of the first dataset at a greater than chance level *(*11). Therefore, for each replication dataset and each module we computed the median of its topological overlap matrix and compared this value against the null distribution of medians computed on random modules of identical size. We used 10,000 re-samplings and finally computed a meta-analysis of module-wise topological replication across the three datasets considered. Additionally, we assessed the concordance of gene loadings in the *ME* across all datasets by means of binomial tests (Figure S2).

**1.2 Module Prioritization**

*Overrepresentation of miR-137 target genes in modules.* We used four miRNA target repositories to obtain lists of predicted miR-137 targets (TargetScan v7.1, <http://www.targetscan.org/vert_71/> *(*12); MirTarget, <http://www.mirdb.org/> *(*13); TargetMiner, <http://www.isical.ac.in/~bioinfo_miu/targetminer20.htm> *(*14), and TarBase V7.0 *(*15)). Then, we performed a hypergeometric test for over-representation of miR-137 target lists in each co-expression module and combined p-values (sum-log Fisher’s method; α_Bonferroni_ = 0.05/(N_replicated_modules_×4)).

*Overrepresentation of SCZ risk loci genes in miR-137 enriched modules.* We assessed the overrepresentation of SCZ risk genes *(*16) (N = 320; list in Table S2), using hypergeometric tests (α_Bonferroni_ = 0.05/N_mod_miR-137_). We validated this measure of SCZ enrichment using MAGMA *(*17) to investigate the overrepresentation of genetic risk variants (rather than just genes) in the modules selected.

**1.3 Experimental Validation and Data Analysis**

*Cell culture and transfections:* Mouse neuroblastoma Neuro-2a cells were purchased from American Type Culture Collection (ATCC). Neuro-2a cells were cultured in Dulbecco’s Modified Eagle Media (DMEM) cell culture medium supplemented with 10% fetal bovine serum, L-glutamine, sodium pyruvate, and penicillin/streptomycin (HyClone) and maintained at 37°C in a humidified chamber supplemented with 5% CO_2_, as per manufacturer’s instructions. PEI (Polyethylenemine 25 kDa linear, Polyscience) transfection reagent was prepared at concentration of 1 mg/ml and used to transiently transfect Neuro-2a cells, with a mass ratio of PEI to DNA of 3:1.

*Plasmids:* A PcDNA3.2/V5 mmu-mir-137 plasmid expressing miR137 courtesy of David Bartel (Addgene plasmid # 26327)*(*18) and a control PcDNA3.2 were used for miR-137 overexpression (OE) conditions. A CRISPR-Cas9 approach*(*19) was used to generate cells that do not express miR-137. sgRNAs against miR-137 were designed using <http://benchling.com/>. Guides were cloned into a puromycin resistance encoding CRISPR-Cas9 vector PX459 V2.0*(*19), obtained from Dr. Feng Zhang’s lab through Addgene (#62988). A U6 primer (5'GAGGGCCTATTTCCCATGATTCC3') was used for cloning validation of CRISPR-Cas9 plasmids by Sanger sequencing.

*Validation of Cas9-induced miR-137-micro-deletions*: To assess mutation efficiency of the CRISPR-Cas9 system, 50–70% confluent Neuro-2a cells were transfected all in one px459 based constructs. To select only transfected cells, 48 hours after transfection cells were incubated with 3µM puromycin for 72 hours followed by 48 hours incubation without puromycin. Cells were lysed using (Tris pH=8.0 0.1M, NaCl 0.2M, EDTA 5mM, SDS 0.4% and proteinase K 0.2mg/ml) buffer, and DNA was precipitated with isopropanol. Phusion Hot Start II DNA Polymerase standard protocol was used to amplify a 592 bp region containing the miR-137 precursor locus using the following primers: forward: 5'TCTAAAGCGGTCTGGGTCAC3', reverse: 5'TGGGTGACCACCAGGTAAAC3'. PCR product were sequenced and TIDE analyses *(*20) of Sanger sequencing was performed to assess mutation efficiency around targeted sites for each sgRNAs (<https://tide.nki.nl/>). Two sgRNAs (guide 1 and guide 3) induced high level of indels inside the targeted genomic loci (g1 – 75%, g3– 96.3%). The level of decomposition confirmed the quality of analysis (Figure S5B, for guide 3). Guide 1 (5’GTATTCTTGGGTGGATAATA3’) targeted the Dicer processing cite of the miR-137 precursor while Guide 3 (5’CTTAAGAATACGCGTAGTCG3’) targeted the Drosha processing site (Figure S5A).

*Validation of Cas9-induced miR-137-KO*: We measured relative quantity of mature miR-137 using qPCR in order to validate impact on miR-137 expression following transfection of px459 based constructs. RNA was isolated with TRI Reagent by Zymo Research and Direct-zol column based miniprep kit. Reverse transcription reaction was performed using TaqMan™ MicroRNA Reverse Transcription Kit and RT primers that are included with the TaqMan microRNA assays for U6 snRNA (TM: 001973) and for mmu-miR-137 (TM: 001129). qPCR was performed using TaqMan probes that are included in the assays mentioned above and TaqMan™ Universal Master Mix II (no UNG) on QuantStudio3 (Applied Biosystems). The results of qPCR for CRISPR-Cas9 mutants revealed no difference for Guide 1 as compared to control while for Guide 3 a massive reduction of mature mmu-mir-137-3p was detected (Figure S5C).

*Titration of miR-137 expression*: Titration of miR-137 expression was conducted over two batches of experiments each containing a shared OE experimental condition to evaluate replicability as listed below:

*Batch 1*

- PX459 V2.0 empty vector (Control for CRISPR-KO condition)

- miR-137 KO cells

- miR OE_.75_ (.75 µg of transfected PcDNA3.2/V5 mmu-mir-137 vector /well)

*Batch 2*

- PcDNA3.2 empty vector (Control for OE condition)

- miR-137 OE_.15_ (.15µg of transfected PcDNA3.2/V5 mmu-mir-137 vector /well)

- miR OE OE_.75b_ (.75 µg of transfected PcDNA3.2/V5 mmu-mir-137 vector /well)

qPCR analysis from separate biological replicates showed similar levels of miR-137 expression in the two control conditions. The OE_.15_ condition resulted in a 15-fold increase of expression from average control conditions. Both OE_.75_ condition led to comparable 55-50 fold increased in miR-137 expression (Table S5).

*Microarray analysis:* Genome wide impact of miR-137 titration was analyzed using microarrays (Mouse Gene 2.0 ST microarrays, Affymetrix). Primary analysis from each conditions (n=3 biological replicates per conditions) in both batches. Microarray analysis were performed by CHU de Québec Research Center (CHUL) Gene Expression Platform, Quebec, Canada.

Microarrays transcriptome-wide gene expression profile served to generate a gene expression matrix, which we normalized for further analysis by correcting expression intensity values with the *affxparser* package in R *(*21). The neuroblastoma cell line dataset included 18 observations across two batches and 12,867 genes. For each gene, we standardized the individual gene expression to the mean and standard deviation of controls separately by batch. This way, gene expression in the experimental condition was transformed as a deviation from the batch-specific control condition for each gene. Then, we pooled the two experimental batches removing control samples to obtain results independent of the data we used to adjust batch effects. In the remaining 12 samples we used the *empiricalBayes* function in the R *WGCNA* package (4) to reduce residual batch effects. This function allows to protect the influence of a condition of interest while removing the effect of confounding variables. Here, we used batch as confounding variable and miR-137 quantification as condition of interest. We assessed dose-dependent gene expression by means of a linear model including the KO (3 samples), OE_.15_ (3 samples), and OE_.75_ (6 samples) conditions (*limma* package *(*22)). We corrected results for multiple comparisons using False Discovery Rate (FDR) q < .05 and thus obtained a list of dose-dependent expressed genes (DDEGs). Finally, we tested the overrepresentation of these mouse cell line DDEGs in the human modules selected for miR-137 target enrichment by means of hypergeometric tests.

**1.4 Polygenic Co-expression Index.**

We aimed to control for ancestry stratification in the SNP identification to compute the PCI. To this purpose, used the *SNPRelate* R Bioconductor package to select a set of indicative autosomal SNPs with MAF > .05 and performed LD pruning using a sliding window of 500 kbp and a pair-wise R^2^< 0.1. Then, we computed genomic eigenvariates and used these variables as nuisance covariates in the following analyses *(*23).

For the module eigengene association analysis, we detected SNPs within each module gene and its 100 kbp up- and down-stream flankers *(10,* 24). We selected SNPs with within-cohort MAF ≥ 0.15 because the sample size was too limited to investigate uncommon variants and pooled minor allele carriers whenever MAF ≤ 0.2 to avoid biasing estimations of population variance with small genotypic groups. Thus, we selected 66,373 SNPs. To prune the SNP list and obtain independent markers of co-expression, we evaluated pair-wise R^2^ between close SNPs (within 250 kbp; LD threshold was R^2^ >0.1 *(*16)). We then iteratively discarded the SNPs with the weaker associations *(8, 10,* 23) to select 1,792 statistically independent SNPs.

We employed a previously published procedure based on Signal Detection Theory to assign weights (A’) to each SNP genotype (8). For each genotypic population of each of the 50 top-ranked SNPs, we computed the A’ weights such that they were positively correlated with the module eigengene. We defined the PCI as the average of A’ values corresponding to all the genotypes included for each subject. In this way, the PCI could be interpreted as the genetically indexed inter-individual variability associated with gene co-expression approximated by the module eigengene.

A relevant issue is how many SNPs need to be included in the PCI. Including too few SNPs may not allow for sufficient predictive power, while too many SNPs may yield overfitting effects which would decrease replication probability in independent datasets. To identify a SNP ensemble with significant predictive power we used BrainEAC as independent replication dataset. We ranked SNPs based on their association with the module eigengene of *Darkorange* and computed 50 PCIs with an increasing number of SNPs (the first PCI included just the first ranked co-eQTL, the second PCI included the first and the second co-eQTL, and so on up to the 50^th^ co-eQTL). We then assessed the Spearman’s correlations between the PCIs and the module eingengene of *Darkorange* computed by projecting module eigengene gene weights onto the BrainEAC data (one-tailed p-value < 0.05). We used the A’ weights derived from the CMC datasets to compute the PCIs in BrainEAC.

**1.5 Neuroimaging study**

*Participants.* All subjects were Caucasians from the region of Apulia, Italy. Inclusion criteria were the absence of any lifetime psychiatric disorder, as evaluated with the Structured Clinical Interview for Diagnostic and Statistical Manual of Mental Disorders IV, NP (Non atient version), of any significant neurological or medical condition revealed by clinical and magnetic resonance imaging evaluation, of history of head trauma with loss of consciousness, and of pharmacological treatment or drug abuse in the past year. The Wechsler Adult Intelligence Scale–Revised was used to evaluate the intelligence quotient *(*25), the Hollingshead Scale *(*26) to calculate the socio-economical status, and the Edinburgh Inventory *(*27) to measure handedness. The present experimental protocol was approved by the local institutional review board.

*EP task.* This task consists of a presentation of angry, fearful, happy and neutral facial expressions from a validated set of facial pictures (NimStim, http://www.macbrain.org/resources.htm). Stimuli order was pseudorandom. From stimulus appearance, 2 s were allowed for behavioral responses. Each stimulus was presented for 500 ms, with a response deadline of 2000 ms from onset. The interstimulus interval pseudo-randomly jittered between 2 and 7 s (inter-stimulus-interval average after jittering: 2.7 s). The total number of stimuli was 144: 30 angry, 39 fearful, 37 happy, and 38 neutral faces. The duration of the task was 6 min 8 s. A fixation crosshair was presented during the interstimulus interval. Subjects indicated whether the face presented was “male” or “female”.

*fMRI data acquisition and analysis.* Blood Oxygen Level Dependent (BOLD) fMRI was performed on a GE Signa 3T scanner (repetition time, 2000 ms; echo time, 28 ms; 26 interleaved axial slices; thickness, 4 mm; gap, 1 mm; voxel size, 3.75 × 3.75 × 5; flip angle, 90°; field of view, 24 cm; matrix, 64 × 64) while participants performed the task. The first four scans were discarded to allow for signal saturation. Stimuli were presented via a back-projection system and responses were recorded through a fiber optic response box which allowed recording behavioral data as percent of correct responses and reaction times. fMRI responses were modeled using a canonical hemodynamic response function and temporally filtered using a high-pass filter of 128 Hz to minimize scanner drift.

Analysis of the fMRI data was completed using Statistical Parametric Mapping 12 (SPM12, Wellcome Department of Cognitive Neurology, London, UK). Images for each subject were realigned to the first volume in the time series and movement parameters were extracted to exclude subjects with excessive head motion (>3.75 mm of translation, >3.75° rotation). We excluded subjects if just one volume in the fMRI time series exceeded the movement threshold. Images were slice timing corrected, re-sampled to a 3.75 mm isotropic voxel size, spatially normalized into a standard stereotactic space (Montreal Institute on Neurology, MNI, template) and smoothed using a 8 mm full-width half-maximum isotropic Gaussian kernel. A box-car model convolved with the hemodynamic response function (HRF) at each voxel was then fitted on the data. Six motion parameters (three translation and three rotation parameters) of the current volume and the preceding volume, obtained from the realignment procedure, plus each of these 12 values squared, were included in the general linear model (GLM) as covariates of no interest *(*28) (Friston 24-parameter model). In the first-level analysis, linear contrasts were computed producing t statistical maps at each voxel including all the faces stimuli taken together (happy + angry + fearful + neutral) vs. crosshair. All individual contrast images were entered in a second-level random effects analysis as reported in the main text. All analyses were constrained by a task-specific activity mask in order to focus on brain regions significantly more active during the task compared to the baseline (p<0.05).

*Connectivity analysis.* In order to assess the effect of PCI_miR-137_ on prefrontal-amygdala coupling we computed a genetic-physiological interaction connectivity analysis *(*29). Specifically, we extracted the first eigenvariates from the individual time series within the prefrontal cluster associated with the PCI_miR-137_ both in the discovery and replication sample. Then, we entered this new regressor, together with task and movement regressors, into a new general linear model with the amygdala time-series (masked via WFU Pickatlas toolbox), as the regressor of interest. Individual connectivity maps were then entered into a second level regression analysis with the PCI_miR-137_ as regressor of interest and gender, age, squared PCI_miR-137_, and 10 genomic eigenvariates as covariates. For this investigation, we used a statistical threshold of SVC-corrected p < 0.05 (minimum cluster size [k] = 10). Estimates of prefrontal-amygdala coupling were then extracted from significant voxels using MarsBar (http://marsbar.sourceforge.net/) for illustrative purposes.

**1.6 Control imaging genetics analyses**

We computed additional fMRI analyses to address the specificity of miR-137 effects and the relationship between the PCI and the PRS. For all these analyses we set the threshold at p < .001 uncorrected to obtain greater sensitivity. We addressed the following research questions:

1. ***Is the lack of a significant PCI_miR-137_ effect in n-back associated with a limited sample size?*** To address this question, we enlarged the n-back sample including participants without Faces data and reached a sample size of N=481 participants. We attempted to replicate the PRS_miR-137_ effect observed in the main sample and we also tested PCI_miR-137_ as a predictor of n-back related brain activity.
2. ***Are PRS_miR-137_ effects specific or do they overlap with the effect of PRSs unrelated with miR-137?*** To answer this question we computed complementary PRSs at the same thresholds employed for the PRS_miR-137_ (meaning these PRSs included all SCZ variants except those associated with miR-137 genes) and assessed their effects in the main sample of 222 individuals during Faces and n-back.
3. ***Are PRS_miR-137_ effects on fMRI phenotypes related with Darkorange genes?*** We intersected *Darkorange* genes with the list provided by Cosgrove, Harold, Mothersill, Anney, Hill, Bray, Blokland, Petryshen, Wellcome Trust Case Control, Richards, Mantripragada, Owen, O'Donovan, Gill, Corvin, Morris and Donohoe *(*30) and tested PRSs based only on variants within these genes.
4. ***Do PRS_miR-137_ effects on fMRI phenotypes persist without Darkorange genes?*** We computed scores complementary to those mentioned in point 3 and tested them on the fMRI phenotypes considered.
2. **SI Results**

Table S7 reports the findings of the control analyses mentioned in SI 1.6. Here is a brief summary of the findings:

1. The analysis of a sample of 481 n-back scans replicated the results reported in the main text for the second and third PRS threshold in the same positive direction. Also in this larger sample we found no significant effect of PCI_miR-137_.
2. The analysis of the main sample of 222 individuals did not reveal any significant effects of the complementary PRSs non-associated with miR-137 during n-back. Instead, we found a negative effect during performance of the Faces task in a medial temporal cluster including the amygdala.
3. PRS_miR-137_ including only variants harbored in *Darkorange* genes did not show any significant associations with either n-back or Faces PFC activations – possibly also because these scores included few variants.
4. Complementary PRS_miR-137_ not including *Darkorange* genes yielded effects essentially indistinguishable from PRS_miR-137_ effects – again a possible consequence of the paucity of variants excluded.

Overall, these additional analyses show that the effects detected were specific of genes associated with miR-137 and the effects of the PRS_miR-137_ and the PCI_miR-137_ did not depend on each other.

**3. SI Figures**

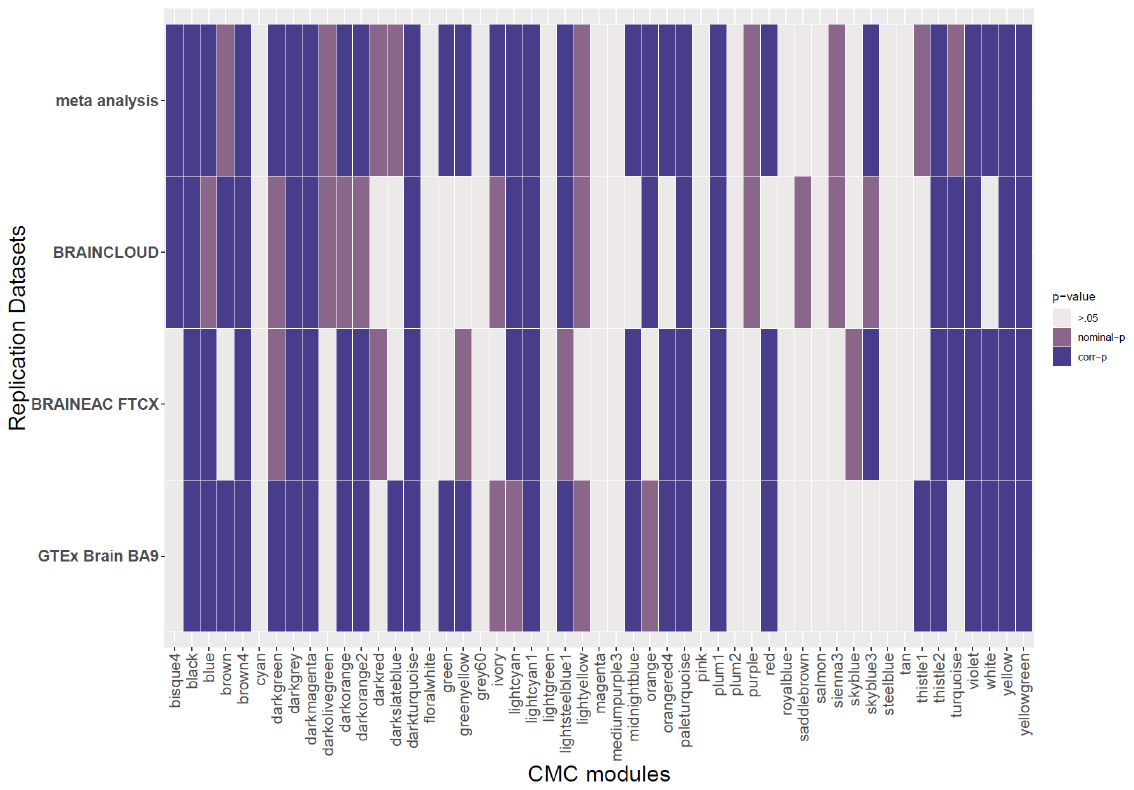

**Figure S1. Module-wise replication.** Color map represents the empirical significance of the topological replication of the CMC modules in three independent datasets. The 28 modules with a purple box in the meta-analysis row were considered for further analyses.

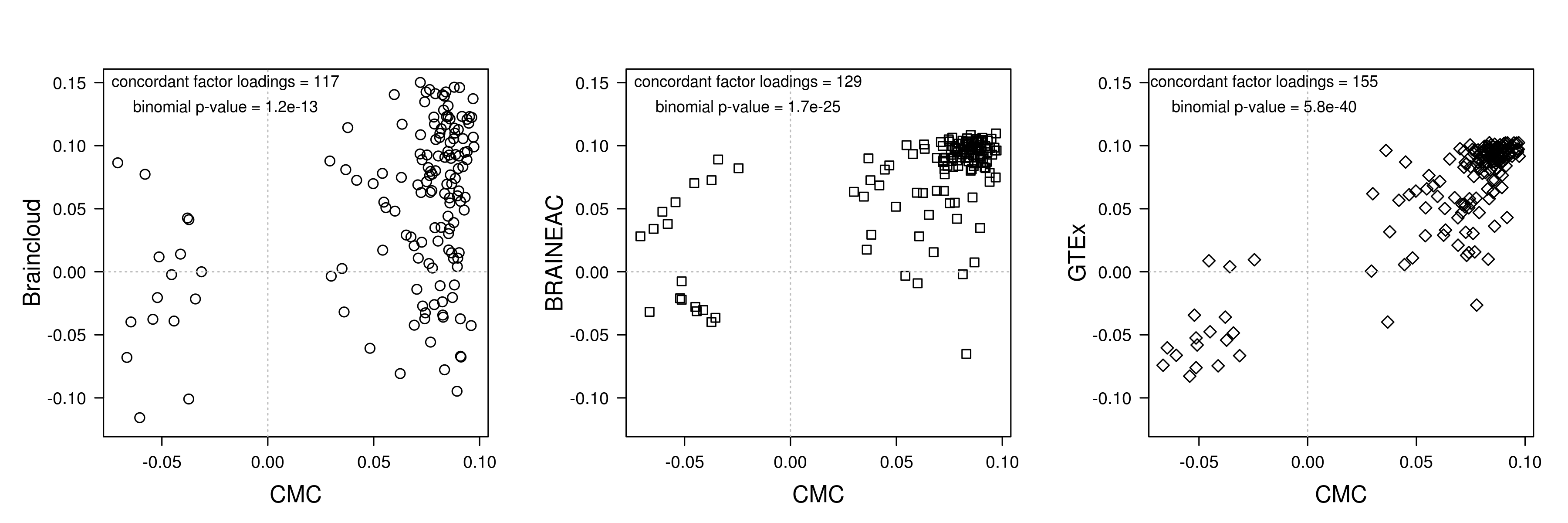

**Figure S2. Gene loading replication on *Darkorange* module eigengene.** Scatterplots show the association between gene loadings across the discovery dataset (CMC) and the three replication datasets. The top right and left bottom quadrants in each plot represent concordant loadings. Binomial tests were computed considering concordant loadings as the successes out of the number of module genes.

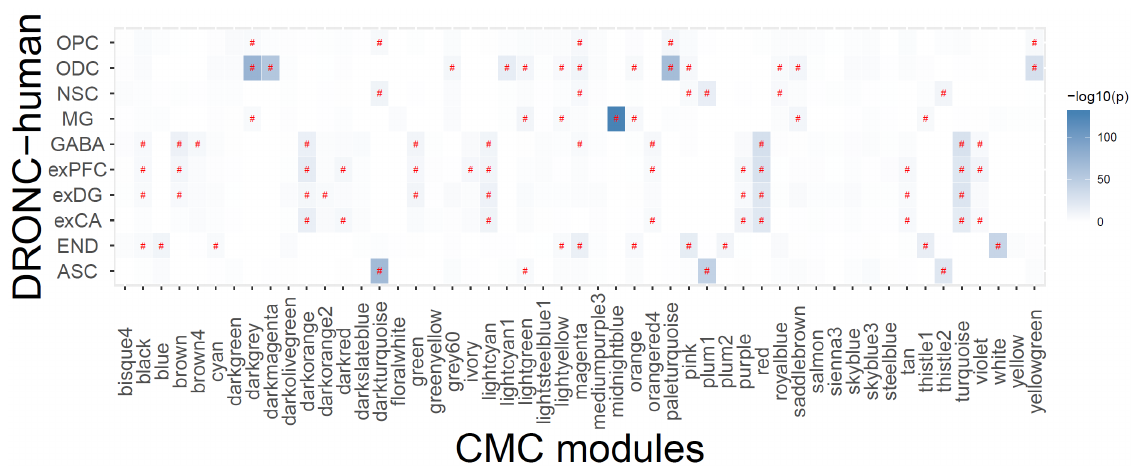

**Figure S3. Module-wise cell specificity.** Color map illustrates the overrepresentation of cell-specific markers within modules. The pound sign denotes significant enrichments after Bonferroni correction for multiple comparisons (α = .05). Abbreviations: ASC=astrocytes, END=endothelial cells, exCA=pyramidal neurons from the Hip CA region, exDG=granule neurons from the hippocampus dentate gyrus region, exPFC=glutamatergic neurons from the PFC, GABA=GABAergic interneurons, MG=microglia, NSC=neuronal stem cells, ODC=oligodendrocytes, OPC=oligodendrocyte precursor cells.

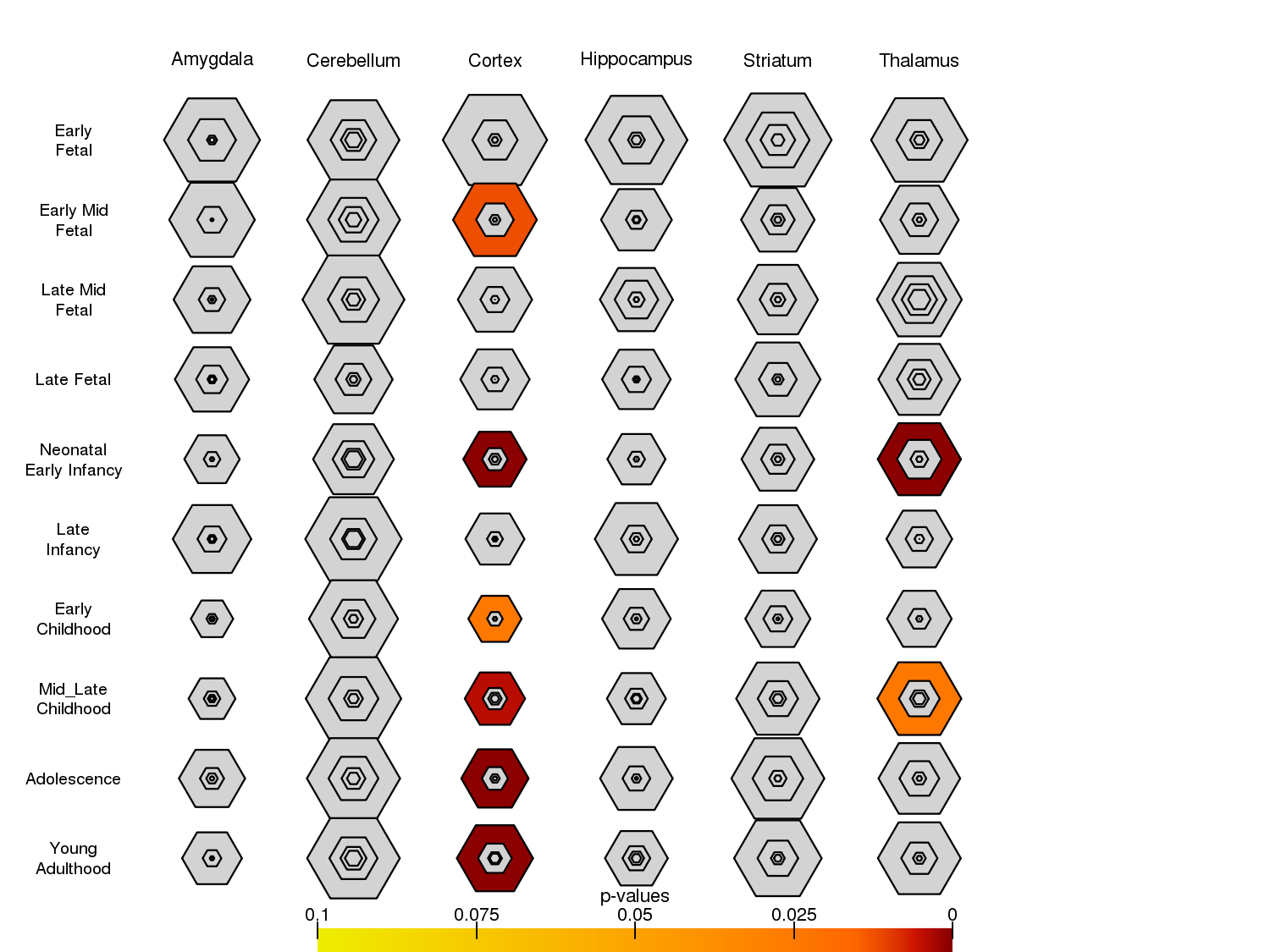

**Figure S4. Brain region specific expression analysis.** The graph shows Darkorange age-specific markers. Colored shapes are overrepresented in the module.

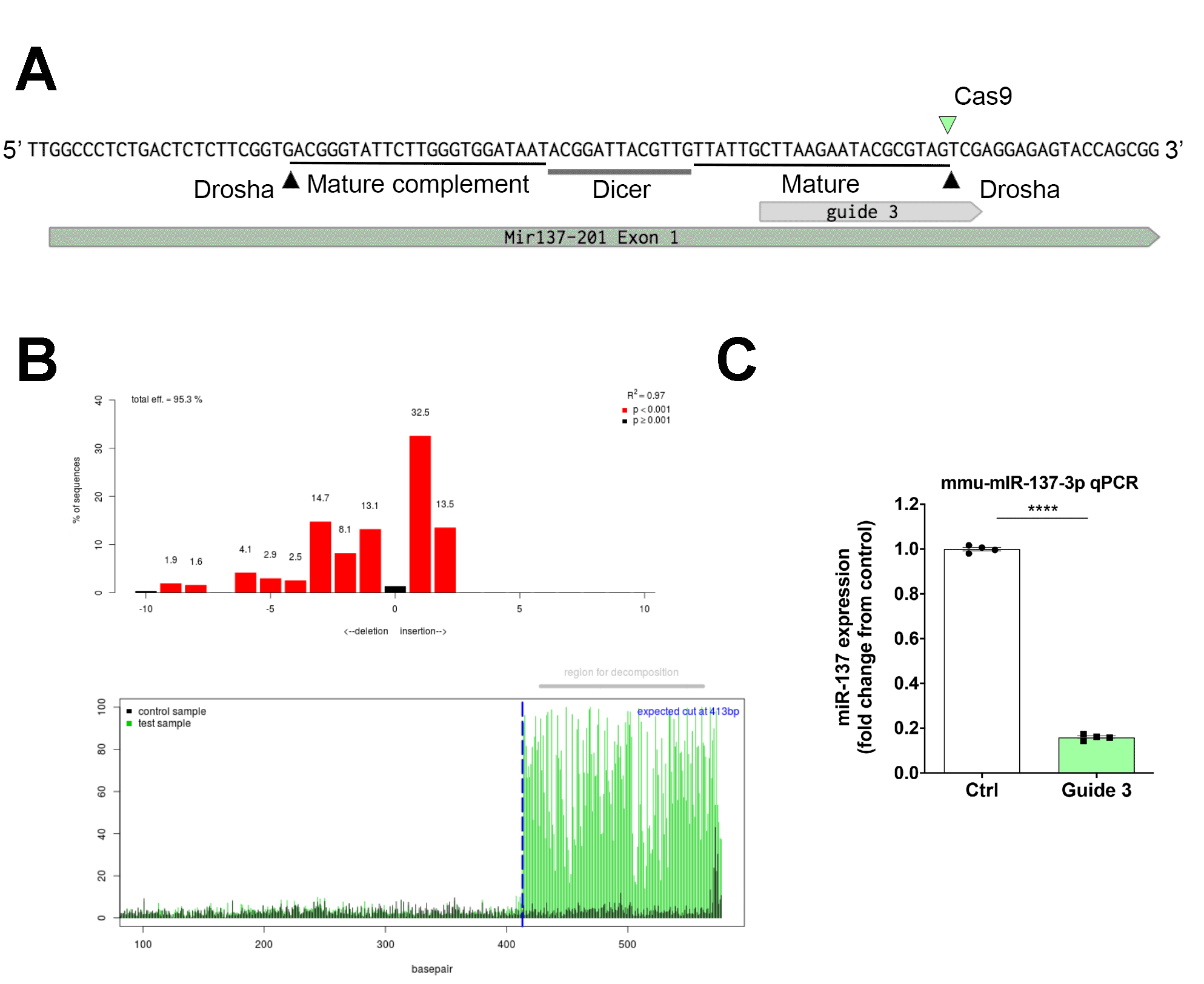

**Figure S5:** Crispr/Cas9 strategy used to knockout down miR137 expression. **A:** Schematic of the miR-137 gene exon 1 encoding the microRNA precursor. Guide 3 led to a Cas9-mediated double strand DNA cut between nucleotides situated at the 3’ terminal of the mature miR-137 sequence. These cuts and internal deletion (indel) repair leads to a disruption of the Drosha processing site of the mutated miR-137 precursor RNA thus preventing the production of mature miR-137. The Cas 9 cutting site (green arrowhead), Drosha cutting sites (black arrowheads), the Dicer targeted precursor loop sequence and the sequence of the mature miR-137 sequence are indicated. **B:** TIDE analyses of CRISPR-CAS9 editing efficiency for guide 3 in cells. (Top panels) Deletion-insertion sequences distribution of genomic DNA for N2A cells after CRISPR-Cas9 modifications. Position “0” demonstrates percentage of non-modified DNA sequences, “+” - insertions, “-” - deletions. (Bottom panels) Percentage of aberrant sequences for mutant and control cells for each guide, blue dashed line marks the cutting site. **C:** Quantification of miR-137 expression using qPCR in N2A neuroblastoma cells transfected with Cas9 and either guide 3 or a control guide RNA. P<0.0001. unpaired Student T-test n=4 separate transfections per group.

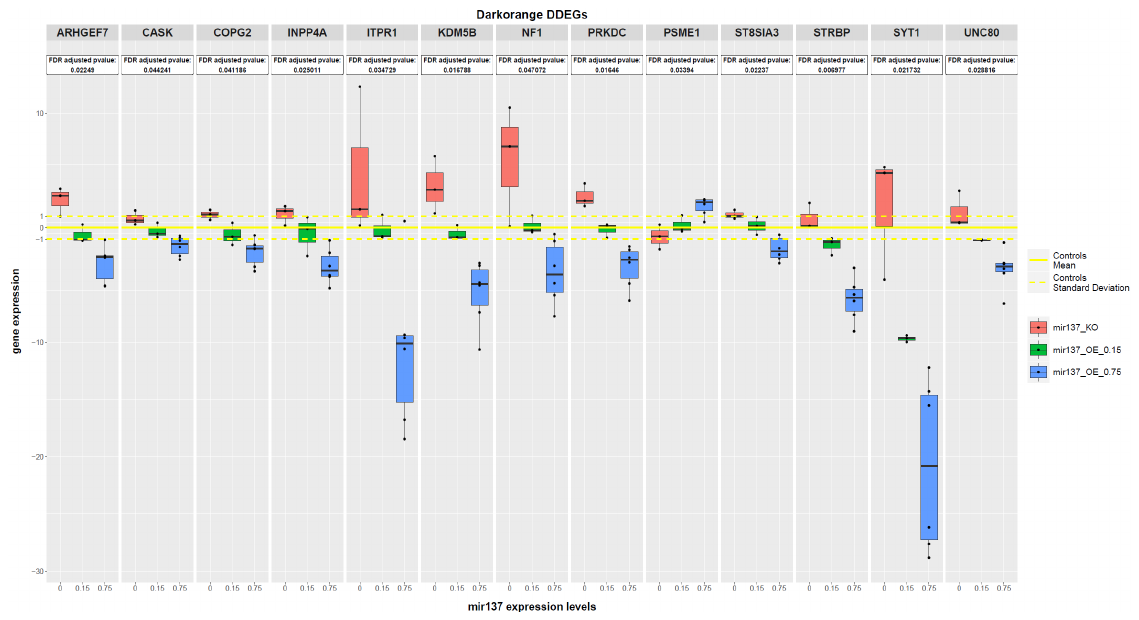

**Figure S6**. **Darkorange dose-dependent expressed genes across experimental conditions.** Boxplots show the expression levels of the 13 Darkorange DDEGs standardized to batch-specific control distribution (mean = 0, standard deviation = 1). Data show that except for gene PSME1 gene expression is negatively correlated with miR-137 expression.

**Figure S7. Association of the Polygenic Co-expression Index with fronto-amygdala connectivity.** Scatter plot depicts the standardized Polygenic Co-expression Index (PCI_miR-137_) on the x-axis and an estimate of connectivity between the replicated dorsolateral prefrontal cluster associated with the PCI_miR-137_ and a bilateral amygdala region of interest.

**4. SI Tables**

**Table S1. Demographics.**

| **Dataset** | **Cohort size** | **Female (male) [ratio]** | **Age mean ± s.d. years** | **Age range (years)** | **IQ mean ± s.d.** | **Handedness mean ± s.d.** | **Socioeconomic status mean ± s.d.** |
| --- | --- | --- | --- | --- | --- | --- | --- |
| CMC | 147 | 54 (93) [0.58] | 61.5 ± 19.9 | 17-95 | - | - | - |
| BrainEAC | 123 | 33 (90) [0.37] | 58 ± 18.7 | 16-102 | - | - | - |
| BrainCloud | 59 | 13 (46) [0.28] | 42.4 ± 16.0 | 17-78 | - | - | - |
| Brainspan | 10 | 4 (6) [0.67] | 7.6 ± 7.9 | 0.3-21 | - | - | - |
| GTEX | 84 | 25 (59) [0.42] | 57.5 ± 10.3 | 23-70 | - | - | - |
| Faces & N-back - Discovery | 222 | 112 (110) [0.50] | 27 ± 7.3 | 18-58 | 109 (11) | 0.7 (0.4) | 42.3 (15.9) |
| Faces - Replication | 136 | 77 (59) [0.57] | 26 ± 7.9 | 18-62 | 104 (13) | 0.7 (0.5) | 35.9 (16.1) |

**Table S2.** Genes in the PGC list included in the network

| **HGNC^a^ symbol** | **Ensemble ID** | **Chromosome** | **Gene**  **Start position** | **Gene**  **End position** | **PGC loci position** | **PGC loci rank** |
| --- | --- | --- | --- | --- | --- | --- |
| SCAND3 | ENSG00000232040 | 6 | 28539407 | 28583989 | chr6:28303247-28712247 | 1 |
| ZKSCAN3 | ENSG00000189298 | 6 | 28317691 | 28336947 | chr6:28303247-28712247 | 1 |
| ZSCAN12 | ENSG00000158691 | 6 | 28346732 | 28367511 | chr6:28303247-28712247 | 1 |
| ZSCAN23 | ENSG00000187987 | 6 | 28399707 | 28411279 | chr6:28303247-28712247 | 1 |
| DPYD | ENSG00000188641 | 1 | 97543299 | 98386605 | chr1:97792625-98559084 | 2 |
| ARL3 | ENSG00000138175 | 10 | 104433488 | 104474164 | chr10:104423800-105165583 | 3 |
| AS3MT | ENSG00000214435 | 10 | 104629273 | 104661656 | chr10:104423800-105165583 | 3 |
| C10orf32 | ENSG00000166275 | 10 | 104613980 | 104624718 | chr10:104423800-105165583 | 3 |
| CNNM2 | ENSG00000148842 | 10 | 104678050 | 104849978 | chr10:104423800-105165583 | 3 |
| INA | ENSG00000148798 | 10 | 105036920 | 105050108 | chr10:104423800-105165583 | 3 |
| NT5C2 | ENSG00000076685 | 10 | 104845940 | 104953056 | chr10:104423800-105165583 | 3 |
| PCGF6 | ENSG00000156374 | 10 | 105062553 | 105110891 | chr10:104423800-105165583 | 3 |
| PDCD11 | ENSG00000148843 | 10 | 105156405 | 105206049 | chr10:104423800-105165583 | 3 |
| SFXN2 | ENSG00000156398 | 10 | 104474295 | 104503249 | chr10:104423800-105165583 | 3 |
| TAF5 | ENSG00000148835 | 10 | 105127724 | 105148822 | chr10:104423800-105165583 | 3 |
| TRIM8 | ENSG00000171206 | 10 | 104404253 | 104418164 | chr10:104423800-105165583 | 3 |
| USMG5 | ENSG00000173915 | 10 | 105148798 | 105156223 | chr10:104423800-105165583 | 3 |
| CACNA1C | ENSG00000151067 | 12 | 2079952 | 2802108 | chr12:2321860-2523731 | 4 |
| TSNARE1 | ENSG00000171045 | 8 | 143293441 | 143484601 | chr8:143309503-143330533 | 5 |
| SLC39A8 | ENSG00000138821 | 4 | 103172198 | 103352415 | chr4:103146888-103198090 | 6 |
| MAD1L1 | ENSG00000002822 | 7 | 1855429 | 2272878 | chr7:1896096-2190096 | 7 |
| ZSWIM6 | ENSG00000130449 | 5 | 60628100 | 60841997 | chr5:60499143-60843543 | 8 |
| ABCB9 | ENSG00000150967 | 12 | 123405498 | 123466196 | chr12:123448113-123909113 | 9 |
| ARL6IP4 | ENSG00000182196 | 12 | 123464607 | 123467456 | chr12:123448113-123909113 | 9 |
| C12orf65 | ENSG00000130921 | 12 | 123717463 | 123742506 | chr12:123448113-123909113 | 9 |
| CDK2AP1 | ENSG00000111328 | 12 | 123745528 | 123756881 | chr12:123448113-123909113 | 9 |
| MPHOSPH9 | ENSG00000051825 | 12 | 123636867 | 123728561 | chr12:123448113-123909113 | 9 |
| OGFOD2 | ENSG00000111325 | 12 | 123459127 | 123464590 | chr12:123448113-123909113 | 9 |
| PITPNM2 | ENSG00000090975 | 12 | 123468027 | 123634562 | chr12:123448113-123909113 | 9 |
| RILPL2 | ENSG00000150977 | 12 | 123899936 | 123921264 | chr12:123448113-123909113 | 9 |
| SBNO1 | ENSG00000139697 | 12 | 123773656 | 123849390 | chr12:123448113-123909113 | 9 |
| SETD8 | ENSG00000183955 | 12 | 123868320 | 123893905 | chr12:123448113-123909113 | 9 |
| C2orf47 | ENSG00000162972 | 2 | 200820040 | 200873263 | chr2:200715237-200848037 | 10 |
| C2orf69 | ENSG00000178074 | 2 | 200775979 | 200820658 | chr2:200715237-200848037 | 10 |
| TYW5 | ENSG00000162971 | 2 | 200794698 | 200820459 | chr2:200715237-200848037 | 10 |
| FES | ENSG00000182511 | 15 | 91426925 | 91439006 | chr15:91416560-91429040 | 11 |
| FURIN | ENSG00000140564 | 15 | 91411822 | 91426688 | chr15:91416560-91429040 | 11 |
| MAN2A2 | ENSG00000196547 | 15 | 91445448 | 91465814 | chr15:91416560-91429040 | 11 |
| TRANK1 | ENSG00000168016 | 3 | 36868311 | 36986548 | chr3:36843183-36945783 | 12 |
| BAG5 | ENSG00000166170 | 14 | 104022881 | 104029168 | chr14:103996234-104184834 | 13 |
| CKB | ENSG00000166165 | 14 | 103985996 | 103989448 | chr14:103996234-104184834 | 13 |
| KLC1 | ENSG00000126214 | 14 | 104028233 | 104167888 | chr14:103996234-104184834 | 13 |
| PPP1R13B | ENSG00000088808 | 14 | 104200089 | 104313927 | chr14:103996234-104184834 | 13 |
| TRMT61A | ENSG00000166166 | 14 | 103995521 | 104003410 | chr14:103996234-104184834 | 13 |
| XRCC3 | ENSG00000126215 | 14 | 104163946 | 104181841 | chr14:103996234-104184834 | 13 |
| ZFYVE21 | ENSG00000100711 | 14 | 104182067 | 104200005 | chr14:103996234-104184834 | 13 |
| CHRNA3 | ENSG00000080644 | 15 | 78885394 | 78913637 | chr15:78803032-78926732 | 14 |
| CHRNA5 | ENSG00000169684 | 15 | 78857862 | 78887611 | chr15:78803032-78926732 | 14 |
| IREB2 | ENSG00000136381 | 15 | 78729773 | 78793798 | chr15:78803032-78926732 | 14 |
| PSMA4 | ENSG00000041357 | 15 | 78832747 | 78841604 | chr15:78803032-78926732 | 14 |
| IMMP2L | ENSG00000184903 | 7 | 110303110 | 111202573 | chr7:110843815-111205915 | 15 |
| SNX19 | ENSG00000120451 | 11 | 130745331 | 130786404 | chr11:130714610-130749330 | 16 |
| ZNF804A | ENSG00000170396 | 2 | 185463093 | 185804219 | chr2:185601420-185785420 | 17 |
| CNKSR2 | ENSG00000149970 | X | 21392536 | 21672813 | chrX:21193266-21570266 | 18 |
| CACNB2 | ENSG00000165995 | 10 | 18429606 | 18830798 | chr10:18681005-18770105 | 19 |
| LRP1 | ENSG00000123384 | 12 | 57522276 | 57607134 | chr12:57428314-57682971 | 20 |
| MYO1A | ENSG00000166866 | 12 | 57422301 | 57444982 | chr12:57428314-57682971 | 20 |
| NAB2 | ENSG00000166886 | 12 | 57482677 | 57489259 | chr12:57428314-57682971 | 20 |
| NDUFA4L2 | ENSG00000185633 | 12 | 57628686 | 57634498 | chr12:57428314-57682971 | 20 |
| NXPH4 | ENSG00000182379 | 12 | 57610578 | 57620232 | chr12:57428314-57682971 | 20 |
| R3HDM2 | ENSG00000179912 | 12 | 57643392 | 57824788 | chr12:57428314-57682971 | 20 |
| SHMT2 | ENSG00000182199 | 12 | 57623110 | 57628718 | chr12:57428314-57682971 | 20 |
| STAC3 | ENSG00000185482 | 12 | 57637236 | 57644976 | chr12:57428314-57682971 | 20 |
| STAT6 | ENSG00000166888 | 12 | 57489191 | 57525922 | chr12:57428314-57682971 | 20 |
| TAC3 | ENSG00000166863 | 12 | 57403784 | 57422667 | chr12:57428314-57682971 | 20 |
| TMEM194A | ENSG00000166881 | 12 | 57449426 | 57481846 | chr12:57428314-57682971 | 20 |
| LRRIQ3 | ENSG00000162620 | 1 | 74491699 | 74663871 | chr1:73766426-73991366 | 21 |
| C2orf82 | ENSG00000182600 | 2 | 233721980 | 233743418 | chr2:233559301-233753501 | 22 |
| EFHD1 | ENSG00000115468 | 2 | 233470767 | 233547491 | chr2:233559301-233753501 | 22 |
| GIGYF2 | ENSG00000204120 | 2 | 233562009 | 233725285 | chr2:233559301-233753501 | 22 |
| KCNJ13 | ENSG00000115474 | 2 | 233631174 | 233641278 | chr2:233559301-233753501 | 22 |
| NGEF | ENSG00000066248 | 2 | 233743396 | 233877982 | chr2:233559301-233753501 | 22 |
| ESAM | ENSG00000149564 | 11 | 124622026 | 124632186 | chr11:124610007-124620147 | 23 |
| NRGN | ENSG00000154146 | 11 | 124609742 | 124617106 | chr11:124610007-124620147 | 23 |
| TCF4 | ENSG00000196628 | 18 | 52889562 | 53332018 | chr18:52747686-53200117 | 24 |
| AMBRA1 | ENSG00000110497 | 11 | 46417964 | 46615675 | chr11:46342943-46751213 | 25 |
| ARHGAP1 | ENSG00000175220 | 11 | 46698630 | 46722165 | chr11:46342943-46751213 | 25 |
| ATG13 | ENSG00000175224 | 11 | 46638826 | 46696368 | chr11:46342943-46751213 | 25 |
| CHRM4 | ENSG00000180720 | 11 | 46406640 | 46408107 | chr11:46342943-46751213 | 25 |
| CKAP5 | ENSG00000175216 | 11 | 46764598 | 46867847 | chr11:46342943-46751213 | 25 |
| CREB3L1 | ENSG00000157613 | 11 | 46299212 | 46342972 | chr11:46342943-46751213 | 25 |
| DGKZ | ENSG00000149091 | 11 | 46354455 | 46402104 | chr11:46342943-46751213 | 25 |
| HARBI1 | ENSG00000180423 | 11 | 46624411 | 46639459 | chr11:46342943-46751213 | 25 |
| MDK | ENSG00000110492 | 11 | 46402306 | 46405375 | chr11:46342943-46751213 | 25 |
| ZNF408 | ENSG00000175213 | 11 | 46722368 | 46727462 | chr11:46342943-46751213 | 25 |
| CCDC39 | ENSG00000145075 | 3 | 180320646 | 180588793 | chr3:180588843-181205585 | 26 |
| DNAJC19 | ENSG00000205981 | 3 | 180701497 | 180707562 | chr3:180588843-181205585 | 26 |
| FXR1 | ENSG00000114416 | 3 | 180585929 | 180700541 | chr3:180588843-181205585 | 26 |
| ACTR5 | ENSG00000101442 | 20 | 37377085 | 37400834 | chr20:37361494-37485994 | 27 |
| PPP1R16B | ENSG00000101445 | 20 | 37434348 | 37551667 | chr20:37361494-37485994 | 27 |
| SLC32A1 | ENSG00000101438 | 20 | 37353105 | 37358015 | chr20:37361494-37485994 | 27 |
| FANCL | ENSG00000115392 | 2 | 58386378 | 58468507 | chr2:57943593-58502192 | 28 |
| VRK2 | ENSG00000028116 | 2 | 58134786 | 58387055 | chr2:57943593-58502192 | 28 |
| ADAMTSL3 | ENSG00000156218 | 15 | 84322838 | 84708594 | chr15:84661161-85153461 | 29 |
| ZSCAN2 | ENSG00000176371 | 15 | 85144217 | 85171027 | chr15:84661161-85153461 | 29 |
| ANKRD44 | ENSG00000065413 | 2 | 197831741 | 198175897 | chr2:198148577-198835577 | 31 |
| BOLL | ENSG00000152430 | 2 | 198591603 | 198651486 | chr2:198148577-198835577 | 31 |
| COQ10B | ENSG00000115520 | 2 | 198318147 | 198340032 | chr2:198148577-198835577 | 31 |
| HSPD1 | ENSG00000144381 | 2 | 198351305 | 198381461 | chr2:198148577-198835577 | 31 |
| HSPE1 | ENSG00000115541 | 2 | 198364718 | 198368181 | chr2:198148577-198835577 | 31 |
| MARS2 | ENSG00000247626 | 2 | 198570087 | 198573113 | chr2:198148577-198835577 | 31 |
| PLCL1 | ENSG00000115896 | 2 | 198669426 | 199437305 | chr2:198148577-198835577 | 31 |
| RFTN2 | ENSG00000162944 | 2 | 198432948 | 198540769 | chr2:198148577-198835577 | 31 |
| SF3B1 | ENSG00000115524 | 2 | 198254508 | 198299815 | chr2:198148577-198835577 | 31 |
| CHADL | ENSG00000100399 | 22 | 41625517 | 41636938 | chr22:41408556-41675156 | 32 |
| EP300 | ENSG00000100393 | 22 | 41487790 | 41576081 | chr22:41408556-41675156 | 32 |
| L3MBTL2 | ENSG00000100395 | 22 | 41601209 | 41627275 | chr22:41408556-41675156 | 32 |
| RANGAP1 | ENSG00000100401 | 22 | 41641615 | 41682255 | chr22:41408556-41675156 | 32 |
| KCNV1 | ENSG00000164794 | 8 | 110975874 | 110988076 | chr8:111460061-111630761 | 33 |
| CNTN4 | ENSG00000144619 | 3 | 2140497 | 3099645 | chr3:2532786-2561686 | 34 |
| DRD2 | ENSG00000149295 | 11 | 113280318 | 113346413 | chr11:113317794-113423994 | 35 |
| IGSF9B | ENSG00000080854 | 11 | 133778459 | 133826880 | chr11:133808069-133852969 | 36 |
| GLT8D1 | ENSG00000016864 | 3 | 52728505 | 52740048 | chr3:52541105-52903405 | 37 |
| GNL3 | ENSG00000163938 | 3 | 52715172 | 52728508 | chr3:52541105-52903405 | 37 |
| ITIH3 | ENSG00000162267 | 3 | 52828784 | 52843025 | chr3:52541105-52903405 | 37 |
| ITIH4 | ENSG00000055955 | 3 | 52846991 | 52865495 | chr3:52541105-52903405 | 37 |
| MUSTN1 | ENSG00000272573 | 3 | 52867130 | 52869235 | chr3:52541105-52903405 | 37 |
| NEK4 | ENSG00000114904 | 3 | 52744800 | 52804965 | chr3:52541105-52903405 | 37 |
| NISCH | ENSG00000010322 | 3 | 52489134 | 52527087 | chr3:52541105-52903405 | 37 |
| NT5DC2 | ENSG00000168268 | 3 | 52558386 | 52569070 | chr3:52541105-52903405 | 37 |
| PBRM1 | ENSG00000163939 | 3 | 52579368 | 52719933 | chr3:52541105-52903405 | 37 |
| SPCS1 | ENSG00000114902 | 3 | 52738971 | 52742182 | chr3:52541105-52903405 | 37 |
| STAB1 | ENSG00000010327 | 3 | 52529354 | 52558511 | chr3:52541105-52903405 | 37 |
| TMEM110 | ENSG00000213533 | 3 | 52870235 | 52931612 | chr3:52541105-52903405 | 37 |
| ALDOA | ENSG00000149925 | 16 | 30064411 | 30081778 | chr16:29924377-30144877 | 38 |
| ASPHD1 | ENSG00000174939 | 16 | 29911696 | 29931185 | chr16:29924377-30144877 | 38 |
| DOC2A | ENSG00000149927 | 16 | 30016830 | 30034591 | chr16:29924377-30144877 | 38 |
| FAM57B | ENSG00000149926 | 16 | 30035748 | 30064299 | chr16:29924377-30144877 | 38 |
| GDPD3 | ENSG00000102886 | 16 | 30116131 | 30125177 | chr16:29924377-30144877 | 38 |
| HIRIP3 | ENSG00000149929 | 16 | 30003645 | 30007757 | chr16:29924377-30144877 | 38 |
| INO80E | ENSG00000169592 | 16 | 30006615 | 30017114 | chr16:29924377-30144877 | 38 |
| KCTD13 | ENSG00000174943 | 16 | 29916333 | 29938356 | chr16:29924377-30144877 | 38 |
| MAPK3 | ENSG00000102882 | 16 | 30125426 | 30134827 | chr16:29924377-30144877 | 38 |
| PPP4C | ENSG00000149923 | 16 | 30087299 | 30096698 | chr16:29924377-30144877 | 38 |
| SEZ6L2 | ENSG00000174938 | 16 | 29882480 | 29910868 | chr16:29924377-30144877 | 38 |
| TAOK2 | ENSG00000149930 | 16 | 29984962 | 30003582 | chr16:29924377-30144877 | 38 |
| TMEM219 | ENSG00000149932 | 16 | 29952206 | 29984373 | chr16:29924377-30144877 | 38 |
| YPEL3 | ENSG00000090238 | 16 | 30103635 | 30108236 | chr16:29924377-30144877 | 38 |
| CACNA1I | ENSG00000100346 | 22 | 39966758 | 40085742 | chr22:39975317-40016817 | 39 |
| MSL2 | ENSG00000174579 | 3 | 135867764 | 135916083 | chr3:135807405-136615405 | 40 |
| NCK1 | ENSG00000158092 | 3 | 136581050 | 136668665 | chr3:135807405-136615405 | 40 |
| PCCB | ENSG00000114054 | 3 | 135969148 | 136056738 | chr3:135807405-136615405 | 40 |
| PPP2R3A | ENSG00000073711 | 3 | 135684515 | 135866733 | chr3:135807405-136615405 | 40 |
| STAG1 | ENSG00000118007 | 3 | 136055077 | 136471220 | chr3:135807405-136615405 | 40 |
| GRIA1 | ENSG00000155511 | 5 | 152869175 | 153193429 | chr5:151941104-152797656 | 41 |
| PJA1 | ENSG00000181191 | X | 68380694 | 68385636 | chrX:68377126-68379036 | 42 |
| SGSM2 | ENSG00000141258 | 17 | 2240792 | 2284352 | chr17:2095899-2220799 | 43 |
| SMG6 | ENSG00000070366 | 17 | 1963133 | 2207065 | chr17:2095899-2220799 | 43 |
| SRR | ENSG00000167720 | 17 | 2206677 | 2228554 | chr17:2095899-2220799 | 43 |
| TSR1 | ENSG00000167721 | 17 | 2225797 | 2240801 | chr17:2095899-2220799 | 43 |
| GRM3 | ENSG00000198822 | 7 | 86273230 | 86494200 | chr7:86403226-86459326 | 44 |
| VPS13C | ENSG00000129003 | 15 | 62144588 | 62352672 | chr15:61831663-61909663 | 45 |
| KDM4A | ENSG00000066135 | 1 | 44115829 | 44171186 | chr1:44029384-44128084 | 46 |
| PTPRF | ENSG00000142949 | 1 | 43990858 | 44089343 | chr1:44029384-44128084 | 46 |
| CILP2 | ENSG00000160161 | 19 | 19649057 | 19657468 | chr19:19374022-19658022 | 47 |
| GATAD2A | ENSG00000167491 | 19 | 19496635 | 19619740 | chr19:19374022-19658022 | 47 |
| HAPLN4 | ENSG00000187664 | 19 | 19366450 | 19373605 | chr19:19374022-19658022 | 47 |
| MAU2 | ENSG00000129933 | 19 | 19431490 | 19469563 | chr19:19374022-19658022 | 47 |
| NCAN | ENSG00000130287 | 19 | 19322782 | 19363042 | chr19:19374022-19658022 | 47 |
| NDUFA13 | ENSG00000186010 | 19 | 19626545 | 19644285 | chr19:19374022-19658022 | 47 |
| PBX4 | ENSG00000105717 | 19 | 19672522 | 19729725 | chr19:19374022-19658022 | 47 |
| SUGP1 | ENSG00000105705 | 19 | 19386827 | 19431653 | chr19:19374022-19658022 | 47 |
| TSSK6 | ENSG00000178093 | 19 | 19623227 | 19626838 | chr19:19374022-19658022 | 47 |
| YJEFN3 | ENSG00000250067 | 19 | 19627036 | 19648390 | chr19:19374022-19658022 | 47 |
| ANP32E | ENSG00000143401 | 1 | 150190717 | 150208504 | chr1:149998890-150242490 | 48 |
| APH1A | ENSG00000117362 | 1 | 150237804 | 150241980 | chr1:149998890-150242490 | 48 |
| C1orf51 | ENSG00000159208 | 1 | 150254953 | 150259505 | chr1:149998890-150242490 | 48 |
| C1orf54 | ENSG00000118292 | 1 | 150240600 | 150253327 | chr1:149998890-150242490 | 48 |
| CA14 | ENSG00000118298 | 1 | 150229554 | 150237478 | chr1:149998890-150242490 | 48 |
| OTUD7B | ENSG00000163113 | 1 | 149909705 | 149982625 | chr1:149998890-150242490 | 48 |
| PLEKHO1 | ENSG00000023902 | 1 | 150121373 | 150136916 | chr1:149998890-150242490 | 48 |
| VPS45 | ENSG00000136631 | 1 | 150039369 | 150117505 | chr1:149998890-150242490 | 48 |
| SNAP91 | ENSG00000065609 | 6 | 84262599 | 84419410 | chr6:84279922-84407274 | 49 |
| PLCH2 | ENSG00000149527 | 1 | 2357419 | 2436969 | chr1:2372401-2402501 | 50 |
| ERCC4 | ENSG00000175595 | 16 | 14014014 | 14046202 | chr16:13728459-13761359 | 51 |
| PUS7 | ENSG00000091127 | 7 | 105080108 | 105162714 | chr7:104598064-105063064 | 52 |
| SRPK2 | ENSG00000135250 | 7 | 104751151 | 105039755 | chr7:104598064-105063064 | 52 |
| RERE | ENSG00000142599 | 1 | 8412457 | 8877702 | chr1:8411184-8638984 | 53 |
| SLC45A1 | ENSG00000162426 | 1 | 8377886 | 8404227 | chr1:8411184-8638984 | 53 |
| ATP2A2 | ENSG00000174437 | 12 | 110718561 | 110788898 | chr12:110723245-110723245 | 54 |
| C4orf27 | ENSG00000056050 | 4 | 170650616 | 170679104 | chr4:170357552-170646052 | 55 |
| CLCN3 | ENSG00000109572 | 4 | 170533784 | 170644824 | chr4:170357552-170646052 | 55 |
| NEK1 | ENSG00000137601 | 4 | 170314426 | 170533780 | chr4:170357552-170646052 | 55 |
| FUT9 | ENSG00000172461 | 6 | 96463860 | 96663488 | chr6:96459651-96459651 | 56 |
| CENPM | ENSG00000100162 | 22 | 42334725 | 42343168 | chr22:42315744-42689414 | 57 |
| CYP2D6 | ENSG00000100197 | 22 | 42522501 | 42526908 | chr22:42315744-42689414 | 57 |
| FAM109B | ENSG00000177096 | 22 | 42470255 | 42475445 | chr22:42315744-42689414 | 57 |
| NAGA | ENSG00000198951 | 22 | 42454358 | 42466846 | chr22:42315744-42689414 | 57 |
| NDUFA6 | ENSG00000184983 | 22 | 42481529 | 42486959 | chr22:42315744-42689414 | 57 |
| 3-Sep | ENSG00000100167 | 22 | 42372276 | 42394225 | chr22:42315744-42689414 | 57 |
| SHISA8 | ENSG00000234965 | 22 | 42307297 | 42310570 | chr22:42315744-42689414 | 57 |
| SREBF2 | ENSG00000198911 | 22 | 42229109 | 42303312 | chr22:42315744-42689414 | 57 |
| TCF20 | ENSG00000100207 | 22 | 42556019 | 42739622 | chr22:42315744-42689414 | 57 |
| TNFRSF13C | ENSG00000159958 | 22 | 42321045 | 42322822 | chr22:42315744-42689414 | 57 |
| WBP2NL | ENSG00000183066 | 22 | 42394729 | 42454460 | chr22:42315744-42689414 | 57 |
| BTBD18 | ENSG00000233436 | 11 | 57510986 | 57519253 | chr11:57386294-57682294 | 59 |
| C11orf31 | ENSG00000211450 | 11 | 57508825 | 57510986 | chr11:57386294-57682294 | 59 |
| CLP1 | ENSG00000172409 | 11 | 57416465 | 57429340 | chr11:57386294-57682294 | 59 |
| CTNND1 | ENSG00000198561 | 11 | 57520715 | 57587018 | chr11:57386294-57682294 | 59 |
| MED19 | ENSG00000156603 | 11 | 57471186 | 57479693 | chr11:57386294-57682294 | 59 |
| SERPING1 | ENSG00000149131 | 11 | 57364860 | 57382326 | chr11:57386294-57682294 | 59 |
| TMX2 | ENSG00000213593 | 11 | 57480072 | 57508445 | chr11:57386294-57682294 | 59 |
| YPEL4 | ENSG00000166793 | 11 | 57412560 | 57417417 | chr11:57386294-57682294 | 59 |
| ZDHHC5 | ENSG00000156599 | 11 | 57435219 | 57468659 | chr11:57386294-57682294 | 59 |
| LUZP2 | ENSG00000187398 | 11 | 24518516 | 25104150 | chr11:24367320-24412990 | 60 |
| DGKI | ENSG00000157680 | 7 | 137065783 | 137531838 | chr7:137039644-137085244 | 62 |
| PTN | ENSG00000105894 | 7 | 136912088 | 137028611 | chr7:137039644-137085244 | 62 |
| TLE1 | ENSG00000196781 | 9 | 84198598 | 84304220 | chr9:84630941-84813641 | 63 |
| AKT3 | ENSG00000117020 | 1 | 243651535 | 244014381 | chr1:243503719-244002945 | 64 |
| SDCCAG8 | ENSG00000054282 | 1 | 243419320 | 243663394 | chr1:243503719-244002945 | 64 |
| ANKRD63 | ENSG00000230778 | 15 | 40573645 | 40574787 | chr15:40566759-40602237 | 65 |
| PAK6 | ENSG00000137843 | 15 | 40509629 | 40569688 | chr15:40566759-40602237 | 65 |
| PLCB2 | ENSG00000137841 | 15 | 40570377 | 40600136 | chr15:40566759-40602237 | 65 |
| ZNF536 | ENSG00000198597 | 19 | 30719197 | 31204445 | chr19:30981643-31039023 | 66 |
| MEF2C | ENSG00000081189 | 5 | 88013975 | 88199922 | chr5:88581331-88854331 | 67 |
| TBC1D5 | ENSG00000131374 | 3 | 17198654 | 18486309 | chr3:17221366-17888266 | 68 |
| CDC25C | ENSG00000158402 | 5 | 137620954 | 137674044 | chr5:137598121-137948092 | 69 |
| CTNNA1 | ENSG00000044115 | 5 | 137946656 | 138270723 | chr5:137598121-137948092 | 69 |
| EGR1 | ENSG00000120738 | 5 | 137801179 | 137805004 | chr5:137598121-137948092 | 69 |
| ETF1 | ENSG00000120705 | 5 | 137841784 | 137878989 | chr5:137598121-137948092 | 69 |
| FAM53C | ENSG00000120709 | 5 | 137667624 | 137685416 | chr5:137598121-137948092 | 69 |
| HSPA9 | ENSG00000113013 | 5 | 137890571 | 137911133 | chr5:137598121-137948092 | 69 |
| KDM3B | ENSG00000120733 | 5 | 137688285 | 137772717 | chr5:137598121-137948092 | 69 |
| REEP2 | ENSG00000132563 | 5 | 137774706 | 137782658 | chr5:137598121-137948092 | 69 |
| BCL11B | ENSG00000127152 | 14 | 99635624 | 99737861 | chr14:99707919-99719219 | 70 |
| RGS6 | ENSG00000182732 | 14 | 72399156 | 73030654 | chr14:72417326-72450526 | 71 |
| HCN1 | ENSG00000164588 | 5 | 45259349 | 45696253 | chr5:45291475-45393775 | 72 |
| CA8 | ENSG00000178538 | 8 | 61099906 | 61193971 | chr8:60475469-60954469 | 73 |
| CYP26B1 | ENSG00000003137 | 2 | 72356367 | 72375167 | chr2:72357335-72368185 | 74 |
| GRAMD1B | ENSG00000023171 | 11 | 123396344 | 123498482 | chr11:123394636-123395986 | 75 |
| SATB2 | ENSG00000119042 | 2 | 200134223 | 200335989 | chr2:200161422-200309252 | 76 |
| GPM6A | ENSG00000150625 | 4 | 176554085 | 176923815 | chr4:176851001-176875801 | 78 |
| CSMD1 | ENSG00000183117 | 8 | 2792875 | 4852494 | chr8:4177794-4192544 | 79 |
| CUL3 | ENSG00000036257 | 2 | 225334867 | 225450110 | chr2:225334096-225467796 | 80 |
| MMP16 | ENSG00000156103 | 8 | 89044237 | 89340254 | chr8:89340626-89753626 | 81 |
| GRIN2A | ENSG00000183454 | 16 | 9852376 | 10276611 | chr16:9875519-9970219 | 82 |
| PRKD1 | ENSG00000184304 | 14 | 30045687 | 30661104 | chr14:30189985-30190316 | 83 |
| ATXN7 | ENSG00000163635 | 3 | 63850233 | 63989138 | chr3:63792650-64004050 | 84 |
| C3orf49 | ENSG00000163632 | 3 | 63805038 | 63834312 | chr3:63792650-64004050 | 84 |
| PSMD6 | ENSG00000163636 | 3 | 63996225 | 64009658 | chr3:63792650-64004050 | 84 |
| THOC7 | ENSG00000163634 | 3 | 63819546 | 63849579 | chr3:63792650-64004050 | 84 |
| ACD | ENSG00000102977 | 16 | 67691415 | 67694713 | chr16:67709340-68311340 | 85 |
| C16orf86 | ENSG00000159761 | 16 | 67700719 | 67702661 | chr16:67709340-68311340 | 85 |
| CENPT | ENSG00000102901 | 16 | 67862060 | 67881714 | chr16:67709340-68311340 | 85 |
| CTRL | ENSG00000141086 | 16 | 67961543 | 67966317 | chr16:67709340-68311340 | 85 |
| DDX28 | ENSG00000182810 | 16 | 68055179 | 68057770 | chr16:67709340-68311340 | 85 |
| DPEP2 | ENSG00000167261 | 16 | 68021297 | 68034489 | chr16:67709340-68311340 | 85 |
| EDC4 | ENSG00000038358 | 16 | 67906926 | 67918406 | chr16:67709340-68311340 | 85 |
| ESRP2 | ENSG00000103067 | 16 | 68263014 | 68272005 | chr16:67709340-68311340 | 85 |
| GFOD2 | ENSG00000141098 | 16 | 67708434 | 67753324 | chr16:67709340-68311340 | 85 |
| LCAT | ENSG00000213398 | 16 | 67973653 | 67978034 | chr16:67709340-68311340 | 85 |
| NFATC3 | ENSG00000072736 | 16 | 68118654 | 68263162 | chr16:67709340-68311340 | 85 |
| NRN1L | ENSG00000188038 | 16 | 67918708 | 67922758 | chr16:67709340-68311340 | 85 |
| NUTF2 | ENSG00000102898 | 16 | 67880635 | 67906470 | chr16:67709340-68311340 | 85 |
| PARD6A | ENSG00000102981 | 16 | 67694849 | 67696681 | chr16:67709340-68311340 | 85 |
| PLA2G15 | ENSG00000103066 | 16 | 68279207 | 68294961 | chr16:67709340-68311340 | 85 |
| PSKH1 | ENSG00000159792 | 16 | 67927175 | 67963581 | chr16:67709340-68311340 | 85 |
| PSMB10 | ENSG00000205220 | 16 | 67968405 | 67970990 | chr16:67709340-68311340 | 85 |
| RANBP10 | ENSG00000141084 | 16 | 67757005 | 67840555 | chr16:67709340-68311340 | 85 |
| SLC12A4 | ENSG00000124067 | 16 | 67977377 | 68003504 | chr16:67709340-68311340 | 85 |
| SLC7A6 | ENSG00000103064 | 16 | 68298433 | 68335722 | chr16:67709340-68311340 | 85 |
| SLC7A6OS | ENSG00000103061 | 16 | 68318406 | 68344849 | chr16:67709340-68311340 | 85 |
| THAP11 | ENSG00000168286 | 16 | 67876213 | 67878097 | chr16:67709340-68311340 | 85 |
| TSNAXIP1 | ENSG00000102904 | 16 | 67840668 | 67866051 | chr16:67709340-68311340 | 85 |
| RLTPR | ENSG00000159753 | 16 | 67678822 | 67691472 | chr16:67709340-68311340 | 85 |
| EPC2 | ENSG00000135999 | 2 | 149402009 | 149545130 | chr2:149390778-149520178 | 86 |
| ATPAF2 | ENSG00000171953 | 17 | 17880723 | 17942523 | chr17:17722402-18030202 | 87 |
| DRG2 | ENSG00000108591 | 17 | 17991200 | 18011285 | chr17:17722402-18030202 | 87 |
| LRRC48 | ENSG00000171962 | 17 | 17876127 | 17920203 | chr17:17722402-18030202 | 87 |
| MYO15A | ENSG00000091536 | 17 | 18012020 | 18083116 | chr17:17722402-18030202 | 87 |
| RAI1 | ENSG00000108557 | 17 | 17584787 | 17714767 | chr17:17722402-18030202 | 87 |
| SREBF1 | ENSG00000072310 | 17 | 17713713 | 17740325 | chr17:17722402-18030202 | 87 |
| TOM1L2 | ENSG00000175662 | 17 | 17746828 | 17875736 | chr17:17722402-18030202 | 87 |
| TLE3 | ENSG00000140332 | 15 | 70340129 | 70390515 | chr15:70573672-70628872 | 88 |
| CNOT1 | ENSG00000125107 | 16 | 58553855 | 58663790 | chr16:58669293-58682833 | 89 |
| SLC38A7 | ENSG00000103042 | 16 | 58699013 | 58719008 | chr16:58669293-58682833 | 89 |
| CLU | ENSG00000120885 | 8 | 27454434 | 27472548 | chr8:27412627-27453627 | 90 |
| EPHX2 | ENSG00000120915 | 8 | 27348296 | 27403081 | chr8:27412627-27453627 | 90 |
| NLGN4X | ENSG00000146938 | X | 5758678 | 6146904 | chrX:5916533-6032733 | 91 |
| RIMS1 | ENSG00000079841 | 6 | 72596406 | 73112845 | chr6:73132701-73171901 | 92 |
| DFNA5 | ENSG00000105928 | 7 | 24737972 | 24809244 | chr7:24619494-24832094 | 93 |
| MPP6 | ENSG00000105926 | 7 | 24612887 | 24733812 | chr7:24619494-24832094 | 93 |
| OSBPL3 | ENSG00000070882 | 7 | 24836158 | 25021253 | chr7:24619494-24832094 | 93 |
| MAN2A1 | ENSG00000112893 | 5 | 109025067 | 109205326 | chr5:109030036-109209066 | 94 |
| PPARGC1A | ENSG00000109819 | 4 | 23756664 | 23905712 | chr4:23366403-23443403 | 95 |
| GALNT10 | ENSG00000164574 | 5 | 153570290 | 153800544 | chr5:153671057-153688217 | 96 |
| C11orf87 | ENSG00000185742 | 11 | 109292846 | 109299840 | chr11:109285471-109610071 | 97 |
| TMTC1 | ENSG00000133687 | 12 | 29653773 | 29937692 | chr12:29905265-29940365 | 99 |
| PODXL | ENSG00000128567 | 7 | 131185021 | 131242976 | chr7:131539263-131567263 | 100 |
| CD46 | ENSG00000117335 | 1 | 207925402 | 207968858 | chr1:207912183-208024083 | 102 |
| KCNB1 | ENSG00000158445 | 20 | 47980414 | 48099184 | chr20:48114136-48131649 | 103 |
| PTGIS | ENSG00000124212 | 20 | 48120411 | 48184683 | chr20:48114136-48131649 | 103 |
| DPP4 | ENSG00000197635 | 2 | 162848751 | 162931052 | chr2:162798555-162910255 | 105 |
| SLC4A10 | ENSG00000144290 | 2 | 162280843 | 162841792 | chr2:162798555-162910255 | 105 |
| NOSIP | ENSG00000142546 | 19 | 50058968 | 50093519 | chr19:50067499-50135399 | 106 |
| PRR12 | ENSG00000126464 | 19 | 50094900 | 50129696 | chr19:50067499-50135399 | 106 |
| PRRG2 | ENSG00000126460 | 19 | 50083903 | 50094272 | chr19:50067499-50135399 | 106 |
| RCN3 | ENSG00000142552 | 19 | 50030875 | 50050219 | chr19:50067499-50135399 | 106 |
| RRAS | ENSG00000126458 | 19 | 50138549 | 50143458 | chr19:50067499-50135399 | 106 |
| SCAF1 | ENSG00000126461 | 19 | 50145382 | 50161899 | chr19:50067499-50135399 | 106 |
| C12orf42 | ENSG00000179088 | 12 | 103631369 | 103889749 | chr12:103559855-103616655 | 107 |
| CD14 | ENSG00000170458 | 5 | 140011313 | 140013286 | chr5:140023664-140222664 | 108 |
| DND1 | ENSG00000256453 | 5 | 140050379 | 140053171 | chr5:140023664-140222664 | 108 |
| HARS | ENSG00000170445 | 5 | 140052758 | 140071609 | chr5:140023664-140222664 | 108 |
| HARS2 | ENSG00000112855 | 5 | 140071011 | 140078889 | chr5:140023664-140222664 | 108 |
| IK | ENSG00000113141 | 5 | 140026643 | 140042064 | chr5:140023664-140222664 | 108 |
| NDUFA2 | ENSG00000131495 | 5 | 140018325 | 140027370 | chr5:140023664-140222664 | 108 |
| PCDHA1 | ENSG00000204970 | 5 | 140165876 | 140391929 | chr5:140023664-140222664 | 108 |
| PCDHA10 | ENSG00000250120 | 5 | 140235595 | 140391929 | chr5:140023664-140222664 | 108 |
| PCDHA2 | ENSG00000204969 | 5 | 140174444 | 140391929 | chr5:140023664-140222664 | 108 |
| PCDHA3 | ENSG00000255408 | 5 | 140180783 | 140391929 | chr5:140023664-140222664 | 108 |
| PCDHA4 | ENSG00000204967 | 5 | 140186659 | 140391929 | chr5:140023664-140222664 | 108 |
| PCDHA5 | ENSG00000204965 | 5 | 140201222 | 140391929 | chr5:140023664-140222664 | 108 |
| PCDHA6 | ENSG00000081842 | 5 | 140207563 | 140391929 | chr5:140023664-140222664 | 108 |
| PCDHA7 | ENSG00000204963 | 5 | 140213969 | 140391929 | chr5:140023664-140222664 | 108 |
| PCDHA8 | ENSG00000204962 | 5 | 140220907 | 140391929 | chr5:140023664-140222664 | 108 |
| PCDHA9 | ENSG00000204961 | 5 | 140227048 | 140391929 | chr5:140023664-140222664 | 108 |
| TMCO6 | ENSG00000113119 | 5 | 140019012 | 140024993 | chr5:140023664-140222664 | 108 |
| WDR55 | ENSG00000120314 | 5 | 140044261 | 140053709 | chr5:140023664-140222664 | 108 |
| ZMAT2 | ENSG00000146007 | 5 | 140078265 | 140086248 | chr5:140023664-140222664 | 108 |

**Table S3. The CMC network: enrichment statistics for schizophrenia loci and mir137 targets**

| **CMC modules** | **Module sizes** | **Mir137 enrichment** | | **PGC enrichment** |
| --- | --- | --- | --- | --- |
|  |  | **Meta-pvalue** | **Bonferroni corrected** | **hyper-geometric**  **pvalue** |
| blue | 910 | 1.18E-16 | 1.32E-14 | 1.70E-01 |
| **darkorange** | **156** | **9.23E-09** | **1.02E-06** | **5.22E-03** |
| lightcyan1 | 76 | 1.86E-07 | 2.05E-05 | 6.23E-02 |
| black | 575 | 1.98E-05 | 0.00215634 | 8.31E-01 |
| darkorange2 | 73 | 7.66E-04 | 0.082755277 | 1.00E+00 |
| plum1 | 98 | 1.53E-03 | 0.163694173 | 1.00E+00 |
| orange | 161 | 6.06E-03 | 0.642772649 | 4.08E-01 |
| yellowgreen | 107 | 6.59E-03 | 0.692450934 | 3.52E-01 |
| lightsteelblue1 | 82 | 1.78E-02 | 1 | 1.00E+00 |
| darkgrey | 161 | 2.14E-02 | 1 | 2.28E-01 |
| violet | 142 | 4.81E-02 | 1 | 9.40E-01 |
| red | 569 | 8.51E-02 | 1 | 1.02E-02 |
| skyblue3 | 103 | 1.38E-01 | 1 | 3.52E-01 |
| greenyellow | 364 | 1.43E-01 | 1 | 3.61E-01 |
| thistle2 | 45 | 1.50E-01 | 1 | 1.00E+00 |
| brown4 | 71 | 1.85E-01 | 1 | 1.00E+00 |
| darkturquoise | 172 | 3.71E-01 | 1 | 4.57E-01 |
| white | 163 | 4.11E-01 | 1 | 1.00E+00 |
| darkgreen | 196 | 6.13E-01 | 1 | 5.37E-01 |
| paleturquoise | 134 | 6.82E-01 | 1 | 7.72E-01 |
| darkmagenta | 127 | 9.22E-01 | 1 | 7.42E-01 |
| yellow | 606 | 9.29E-01 | 1 | 6.90E-01 |
| orangered4 | 91 | 9.43E-01 | 1 | 8.45E-01 |
| midnightblue | 293 | 9.94E-01 | 1 | 9.78E-01 |
| ivory | 68 | 9.94E-01 | 1 | 1.87E-01 |
| bisque4 | 55 | 9.99E-01 | 1 | 2.41E-02 |
| green | 587 | 9.99E-01 | 1 | 7.78E-01 |
| lightcyan | 271 | 1.00E+00 | 1 | 2.91E-01 |

**Table S4. *Darkorange* genes and overlap with PGC, mir137 targets and mir37** DDEGs

| **HGNC symbol** | **Ensembl ID** | **Chr** | **Gene start** | **Gene end** | **Gene biotype** | **mir137** | **TarBase** | **MirTarget** | **TargetScan** | **TargetMiner** | **mir137 DEGs mouse** | **PGC** |
| --- | --- | --- | --- | --- | --- | --- | --- | --- | --- | --- | --- | --- |
| ABI2 | ENSG00000138443 | 2 | 204192942 | 204312446 | protein_coding |  |  |  |  |  |  |  |
| ACP6 | ENSG00000162836 | 1 | 147101453 | 147142618 | protein_coding |  |  |  |  |  |  |  |
| AKAP6 | ENSG00000151320 | 14 | 32798479 | 33300567 | protein_coding |  |  |  |  |  |  |  |
| AKIRIN2 | ENSG00000135334 | 6 | 88384790 | 88411927 | protein_coding |  |  |  |  |  |  |  |
| ALS2 | ENSG00000003393 | 2 | 202565277 | 202645912 | protein_coding |  |  |  |  |  |  |  |
| AMOTL1 | ENSG00000166025 | 11 | 94439597 | 94609918 | protein_coding |  |  |  |  |  |  |  |
| ANAPC1 | ENSG00000153107 | 2 | 112523848 | 112642267 | protein_coding |  |  |  |  |  |  |  |
| ANK2 | ENSG00000145362 | 4 | 113739265 | 114304896 | protein_coding |  |  |  |  |  |  |  |
| ANKRD18DP | ENSG00000226435 | 3 | 198053522 | 198080720 | protein_coding |  |  |  |  |  |  |  |
| ANKRD34C | ENSG00000235711 | 15 | 79575146 | 79590580 | protein_coding |  |  |  |  |  |  |  |
| ARHGAP20 | ENSG00000137727 | 11 | 110447766 | 110583912 | protein_coding |  |  |  |  |  |  |  |
| ARHGAP26 | ENSG00000145819 | 5 | 142149949 | 142608576 | protein_coding |  |  |  |  |  |  |  |
| ARHGEF3 | ENSG00000163947 | 3 | 56761446 | 57113357 | protein_coding |  |  |  |  |  |  |  |
| ARHGEF7 | ENSG00000102606 | 13 | 111766906 | 111958084 | protein_coding | ✓ | ✓ |  |  |  | ✓ |  |
| ASNS | ENSG00000070669 | 7 | 97481430 | 97501854 | protein_coding |  |  |  |  |  |  |  |
| ATP2A2 | ENSG00000174437 | 12 | 110718561 | 110788898 | protein_coding | ✓ |  |  | ✓ |  |  | ✓ |
| BCL11B | ENSG00000127152 | 14 | 99635624 | 99737861 | protein_coding | ✓ |  |  | ✓ |  |  | ✓ |
| BEND4 | ENSG00000188848 | 4 | 42112955 | 42154895 | protein_coding | ✓ |  |  |  | ✓ |  |  |
| BTRC | ENSG00000166167 | 10 | 103113820 | 103317078 | protein_coding |  |  |  |  |  |  |  |
| C1orf173 | ENSG00000178965 | 1 | 75033795 | 75139422 | protein_coding |  |  |  |  |  |  |  |
| C1orf21 | ENSG00000116667 | 1 | 184356192 | 184598154 | protein_coding |  |  |  |  |  |  |  |
| C1orf63 | ENSG00000117616 | 1 | 25568728 | 25664704 | protein_coding |  |  |  |  |  |  |  |
| CACNA2D1 | ENSG00000153956 | 7 | 81575760 | 82073114 | protein_coding |  |  |  |  |  |  |  |
| CACNB2 | ENSG00000165995 | 10 | 18429606 | 18830798 | protein_coding | ✓ |  |  | ✓ | ✓ |  | ✓ |
| CAP2 | ENSG00000112186 | 6 | 17393447 | 17558023 | protein_coding |  |  |  | ✓ | ✓ |  |  |
| CASK | ENSG00000147044 | X | 41374187 | 41782716 | protein_coding |  |  |  |  |  | ✓ |  |
| CASP10 | ENSG00000003400 | 2 | 202047604 | 202094129 | protein_coding |  |  |  |  |  |  |  |
| CDCA4 | ENSG00000170779 | 14 | 105475910 | 105487485 | protein_coding |  |  |  |  |  |  |  |
| CKAP5 | ENSG00000175216 | 11 | 46764598 | 46867847 | protein_coding |  |  |  |  |  |  | ✓ |
| CNOT1 | ENSG00000125107 | 16 | 58553855 | 58663790 | protein_coding |  |  |  |  |  |  | ✓ |
| CNTNAP2 | ENSG00000174469 | 7 | 145813453 | 148118090 | protein_coding | ✓ |  |  |  | ✓ |  |  |
| COPG2 | ENSG00000158623 | 7 | 130146089 | 130353598 | protein_coding |  |  |  |  |  | ✓ |  |
| CTR9 | ENSG00000198730 | 11 | 10772534 | 10801290 | protein_coding |  |  |  |  |  |  |  |
| DCC | ENSG00000187323 | 18 | 49866542 | 51057784 | protein_coding |  |  |  |  |  |  |  |
| DCLK1 | ENSG00000133083 | 13 | 36345478 | 36705443 | protein_coding | ✓ |  |  |  | ✓ |  |  |
| DLG2 | ENSG00000150672 | 11 | 83166055 | 85338966 | protein_coding |  |  |  |  |  |  |  |
| DLGAP1 | ENSG00000170579 | 18 | 3496030 | 4455335 | protein_coding |  |  |  |  |  |  |  |
| DNAJC16 | ENSG00000116138 | 1 | 15853308 | 15918874 | protein_coding |  |  |  |  |  |  |  |
| DNAJC6 | ENSG00000116675 | 1 | 65713902 | 65881552 | protein_coding |  |  |  |  |  |  |  |
| EFNB2 | ENSG00000125266 | 13 | 107142079 | 107187462 | protein_coding | ✓ |  |  | ✓ |  |  |  |
| EHBP1 | ENSG00000115504 | 2 | 62900986 | 63273622 | protein_coding |  |  |  |  |  |  |  |
| EIF2D | ENSG00000143486 | 1 | 206744620 | 206785904 | protein_coding |  |  |  |  |  |  |  |
| EIF4G3 | ENSG00000075151 | 1 | 21132963 | 21503377 | protein_coding |  |  |  |  |  |  |  |
| EMC8 | ENSG00000131148 | 16 | 85805364 | 85833214 | protein_coding |  |  |  |  |  |  |  |
| ETV5 | ENSG00000244405 | 3 | 185764097 | 185828107 | protein_coding | ✓ |  |  | ✓ | ✓ |  |  |
| FAM135B | ENSG00000147724 | 8 | 139142266 | 139509065 | protein_coding | ✓ |  |  | ✓ | ✓ |  |  |
| FAM160A1 | ENSG00000164142 | 4 | 152330368 | 152584784 | protein_coding |  |  |  |  |  |  |  |
| FAM181A-AS1 | ENSG00000258584 | 14 | 94371076 | 94393412 | processed_transcript |  |  |  |  |  |  |  |
| FARSB | ENSG00000116120 | 2 | 223435255 | 223521056 | protein_coding |  |  |  |  |  |  |  |
| FRMPD4 | ENSG00000169933 | X | 12156585 | 12742642 | protein_coding |  |  |  |  |  |  |  |
| GABRB3 | ENSG00000166206 | 15 | 26788693 | 27184686 | protein_coding |  |  |  |  |  |  |  |
| GPC6 | ENSG00000183098 | 13 | 93879095 | 95059655 | protein_coding |  |  |  |  |  |  |  |
| GPR158 | ENSG00000151025 | 10 | 25463991 | 25891155 | protein_coding | ✓ |  |  | ✓ |  |  |  |
| GRIA3 | ENSG00000125675 | X | 122318006 | 122624766 | protein_coding |  |  |  |  |  |  |  |
| GRIN2A | ENSG00000183454 | 16 | 9852376 | 10276611 | protein_coding | ✓ | ✓ |  | ✓ | ✓ |  | ✓ |
| GRM5 | ENSG00000168959 | 11 | 88237744 | 88799113 | protein_coding | ✓ | ✓ |  | ✓ | ✓ |  |  |
| GSK3B | ENSG00000082701 | 3 | 119540170 | 119813264 | protein_coding | ✓ | ✓ |  | ✓ | ✓ |  |  |
| HERC3 | ENSG00000138641 | 4 | 89442199 | 89629693 | protein_coding | ✓ | ✓ |  | ✓ | ✓ |  |  |
| HS6ST3 | ENSG00000185352 | 13 | 96743093 | 97485671 | protein_coding |  |  |  |  |  |  |  |
| HSD17B7P2 | ENSG00000099251 | 10 | 38645305 | 38667433 | pseudogene |  |  |  |  |  |  |  |
| HSPA12A | ENSG00000165868 | 10 | 118430703 | 118502085 | protein_coding |  |  |  |  |  |  |  |
| ILDR2 | ENSG00000143195 | 1 | 166882443 | 166944719 | protein_coding |  |  |  |  |  |  |  |
| INPP4A | ENSG00000040933 | 2 | 99061317 | 99210853 | protein_coding |  |  |  |  |  | ✓ |  |
| IPCEF1 | ENSG00000074706 | 6 | 154475631 | 154677926 | protein_coding |  |  |  |  |  |  |  |
| ITPR1 | ENSG00000150995 | 3 | 4535032 | 4889524 | protein_coding |  |  |  |  |  | ✓ |  |
| KCNA4 | ENSG00000182255 | 11 | 30031288 | 30038570 | protein_coding |  |  |  |  |  |  |  |
| KCNMA1 | ENSG00000156113 | 10 | 78629359 | 79398353 | protein_coding | ✓ |  |  |  | ✓ |  |  |
| KCNQ3 | ENSG00000184156 | 8 | 133133108 | 133493200 | protein_coding |  |  |  |  |  |  |  |
| KCNQ5 | ENSG00000185760 | 6 | 73331520 | 73908574 | protein_coding | ✓ | ✓ |  |  |  |  |  |
| KCNV1 | ENSG00000164794 | 8 | 110975874 | 110988076 | protein_coding |  |  |  |  |  |  | ✓ |
| KDM5B | ENSG00000117139 | 1 | 202696526 | 202778598 | protein_coding | ✓ |  | ✓ | ✓ |  | ✓ |  |
| KIAA1239 | ENSG00000174145 | 4 | 37245842 | 37451087 | protein_coding |  |  |  |  |  |  |  |
| KIAA1279 | ENSG00000198954 | 10 | 70748487 | 70776738 | protein_coding |  |  |  |  |  |  |  |
| KIAA1549L | ENSG00000110427 | 11 | 33563618 | 33695648 | protein_coding |  |  |  |  |  |  |  |
| KIAA2022 | ENSG00000050030 | X | 73952684 | 74145282 | protein_coding |  |  |  |  |  |  |  |
| KIF1B | ENSG00000054523 | 1 | 10270863 | 10441661 | protein_coding |  |  |  |  |  |  |  |
| KPNA6 | ENSG00000025800 | 1 | 32573639 | 32642169 | protein_coding |  |  |  |  |  |  |  |
| LANCL3 | ENSG00000147036 | X | 37430822 | 37543716 | protein_coding |  |  |  |  |  |  |  |
| LCLAT1 | ENSG00000172954 | 2 | 30670092 | 30867091 | protein_coding |  |  |  |  |  |  |  |
| LDLRAD4 | ENSG00000168675 | 18 | 13217497 | 13652754 | protein_coding | ✓ |  |  | ✓ | ✓ | ✓ |  |
| LMAN2L | ENSG00000114988 | 2 | 97371666 | 97405801 | protein_coding |  |  |  |  |  |  |  |
| LMO7 | ENSG00000136153 | 13 | 76194570 | 76434004 | protein_coding | ✓ |  |  |  | ✓ |  |  |
| LONRF2 | ENSG00000170500 | 2 | 100889753 | 100939195 | protein_coding |  |  |  |  |  |  |  |
| MAD2L2 | ENSG00000116670 | 1 | 11734537 | 11751707 | protein_coding |  |  |  |  |  |  |  |
| MAP1B | ENSG00000131711 | 5 | 71403061 | 71505395 | protein_coding | ✓ |  |  |  | ✓ |  |  |
| MAP2 | ENSG00000078018 | 2 | 210288782 | 210598842 | protein_coding |  |  |  |  |  |  |  |
| METAP1 | ENSG00000164024 | 4 | 99916771 | 99983964 | protein_coding | ✓ |  |  |  | ✓ |  |  |
| MFSD6 | ENSG00000151690 | 2 | 191273081 | 191373931 | protein_coding |  |  |  |  |  |  |  |
| MGAT5 | ENSG00000152127 | 2 | 134119983 | 134454621 | protein_coding |  |  |  |  |  |  |  |
| MYCBP2 | ENSG00000005810 | 13 | 77618792 | 77901185 | protein_coding |  |  |  |  |  |  |  |
| MYO5A | ENSG00000197535 | 15 | 52599480 | 52821247 | protein_coding | ✓ |  |  |  | ✓ |  |  |
| MYRIP | ENSG00000170011 | 3 | 39850405 | 40301812 | protein_coding |  |  |  |  |  |  |  |
| NEDD4L | ENSG00000049759 | 18 | 55711599 | 56068772 | protein_coding |  |  |  |  |  |  |  |
| NF1 | ENSG00000196712 | 17 | 29421945 | 29709134 | protein_coding | ✓ | ✓ | ✓ | ✓ | ✓ | ✓ |  |
| NLGN4X | ENSG00000146938 | X | 5758678 | 6146904 | protein_coding | ✓ | ✓ |  | ✓ |  |  | ✓ |
| NR4A2 | ENSG00000153234 | 2 | 157180944 | 157198860 | protein_coding |  |  |  |  |  |  |  |
| NR4A3 | ENSG00000119508 | 9 | 102584137 | 102629173 | protein_coding | ✓ |  |  |  | ✓ |  |  |
| NRCAM | ENSG00000091129 | 7 | 107788068 | 108097161 | protein_coding |  |  |  |  |  |  |  |
| ✓NRG3 | ENSG00000185737 | 10 | 83635070 | 84746935 | protein_coding | ✓ |  | ✓ | ✓ | ✓ |  |  |
| NRSN1 | ENSG00000152954 | 6 | 24126350 | 24155128 | protein_coding |  |  |  |  |  |  |  |
| ✓OPCML | ENSG00000183715 | 11 | 132284871 | 133402414 | protein_coding | ✓ |  |  | ✓ | ✓ | ✓ |  |
| OSBP | ENSG00000110048 | 11 | 59341871 | 59383617 | protein_coding |  |  |  |  |  |  |  |
| OXGR1 | ENSG00000165621 | 13 | 97637973 | 97646984 | protein_coding |  |  |  |  |  |  |  |
| PARM1 | ENSG00000169116 | 4 | 75858305 | 75975325 | protein_coding |  |  |  |  |  |  |  |
| PART1 | ENSG00000152931 | 5 | 59783540 | 59843484 | lincRNA |  |  |  |  |  |  |  |
| PAXIP1-AS1 | ENSG00000273344 | 7 | 154795158 | 154797413 | lincRNA |  |  |  |  |  |  |  |
| PCDH17 | ENSG00000118946 | 13 | 58205944 | 58303445 | protein_coding | ✓ |  |  |  | ✓ |  |  |
| PCDH19 | ENSG00000165194 | X | 99546642 | 99665271 | protein_coding |  |  |  |  |  |  |  |
| PCDHB14 | ENSG00000120327 | 5 | 140602931 | 140605858 | protein_coding |  |  |  |  |  |  |  |
| PDK3 | ENSG00000067992 | X | 24483338 | 24557954 | protein_coding |  |  |  |  |  |  |  |
| PEG3 | ENSG00000198300 | 19 | 57321445 | 57352096 | protein_coding |  |  |  |  |  |  |  |
| ✓PHLPP2 | ENSG00000040199 | 16 | 71671738 | 71758604 | protein_coding | ✓ | ✓ |  | ✓ | ✓ |  |  |
| PLCB1 | ENSG00000182621 | 20 | 8112824 | 8949003 | protein_coding | ✓ | ✓ | ✓ | ✓ | ✓ |  |  |
| PPAPDC2 | ENSG00000205808 | 9 | 4662298 | 4665256 | protein_coding |  |  |  |  |  |  |  |
| PPTC7 | ENSG00000196850 | 12 | 110969120 | 111021125 | protein_coding | ✓ |  |  | ✓ |  |  |  |
| PRDM10 | ENSG00000170325 | 11 | 129769601 | 129872730 | protein_coding |  |  |  |  |  |  |  |
| PRICKLE2 | ENSG00000163637 | 3 | 64079543 | 64431152 | protein_coding | ✓ |  |  |  | ✓ |  |  |
| PRKCB | ENSG00000166501 | 16 | 23847322 | 24231932 | protein_coding |  |  |  |  |  |  |  |
| PRKDC | ENSG00000253729 | 8 | 48685669 | 48872743 | protein_coding |  |  |  |  |  | ✓ |  |
| PSME1 | ENSG00000092010 | 14 | 24605367 | 24608176 | protein_coding |  |  |  |  |  | ✓ |  |
| PTGER4P2-CDK2AP2P2 |  | 9 | 66494269 | 66503030 | transcribed_pseudogene |  |  |  |  |  |  |  |
| PTPRG | ENSG00000144724 | 3 | 61547243 | 62283288 | protein_coding |  |  |  |  |  |  |  |
| PTPRJ | ENSG00000149177 | 11 | 48002113 | 48189670 | protein_coding |  |  |  |  |  |  |  |
| RASAL2 | ENSG00000075391 | 1 | 178062864 | 178448644 | protein_coding | ✓ | ✓ | ✓ | ✓ |  |  |  |
| RBFOX1 | ENSG00000078328 | 16 | 6069095 | 7763340 | protein_coding |  |  |  |  |  |  |  |
| RBFOX2 | ENSG00000100320 | 22 | 36134783 | 36424473 | protein_coding |  |  |  |  |  |  |  |
| RCC1 | ENSG00000180198 | 1 | 28832455 | 28865812 | protein_coding | ✓ |  | ✓ | ✓ | ✓ |  |  |
| REEP1 | ENSG00000068615 | 2 | 86441116 | 86565206 | protein_coding |  |  |  |  |  |  |  |
| RGMB | ENSG00000174136 | 5 | 98104354 | 98134347 | protein_coding | ✓ |  |  |  | ✓ |  |  |
| RTN3 | ENSG00000133318 | 11 | 63448918 | 63527363 | protein_coding |  |  |  |  |  |  |  |
| SDHAF2 | ENSG00000167985 | 11 | 61197514 | 61215001 | protein_coding |  |  |  |  |  |  |  |
| SGK3 | ENSG00000104205 | 8 | 67624653 | 67774257 | protein_coding |  |  |  |  |  |  |  |
| SH3RF1 | ENSG00000154447 | 4 | 170015407 | 170192256 | protein_coding |  |  |  |  |  |  |  |
| SLC4A8 | ENSG00000050438 | 12 | 51785101 | 51902980 | protein_coding |  |  |  |  |  |  |  |
| SNAP91 | ENSG00000065609 | 6 | 84262599 | 84419410 | protein_coding | ✓ | ✓ |  | ✓ |  |  | ✓ |
| SNX27 | ENSG00000143376 | 1 | 151584541 | 151671567 | protein_coding | ✓ | ✓ |  | ✓ |  |  |  |
| SON | ENSG00000159140 | 21 | 34914924 | 34949812 | protein_coding | ✓ | ✓ |  |  |  |  |  |
| SORL1 | ENSG00000137642 | 11 | 121322912 | 121504402 | protein_coding | ✓ | ✓ |  | ✓ |  |  |  |
| ST8SIA3 | ENSG00000177511 | 18 | 55018044 | 55038962 | protein_coding |  |  |  |  |  | ✓ |  |
| STARD13 | ENSG00000133121 | 13 | 33677272 | 33924767 | protein_coding | ✓ | ✓ |  |  | ✓ |  |  |
| STRBP | ENSG00000165209 | 9 | 125871779 | 126030855 | protein_coding | ✓ | ✓ |  | ✓ |  | ✓ |  |
| STS | ENSG00000101846 | X | 7137497 | 7272851 | protein_coding | ✓ | ✓ |  |  | ✓ |  |  |
| SUSD5 | ENSG00000173705 | 3 | 33191537 | 33260707 | protein_coding |  |  |  |  |  |  |  |
| SYNJ1 | ENSG00000159082 | 21 | 34001069 | 34100359 | protein_coding | ✓ |  |  | ✓ | ✓ |  |  |
| SYT1 | ENSG00000067715 | 12 | 79257773 | 79845788 | protein_coding | ✓ |  | ✓ | ✓ |  | ✓ |  |
| SYT16 | ENSG00000139973 | 14 | 62453803 | 62568431 | protein_coding |  |  |  |  |  |  |  |
| TEX2 | ENSG00000136478 | 17 | 62224587 | 62340661 | protein_coding | ✓ |  |  | ✓ |  |  |  |
| TFCP2 | ENSG00000135457 | 12 | 51487446 | 51566926 | protein_coding |  |  |  |  |  |  |  |
| TGOLN2 | ENSG00000152291 | 2 | 85545147 | 85555548 | protein_coding |  |  |  |  |  |  |  |
| TMEM132B | ENSG00000139364 | 12 | 125671382 | 126146917 | protein_coding | ✓ |  | ✓ | ✓ | ✓ |  |  |
| TMEM178B | ENSG00000261115 | 7 | 140774032 | 141180180 | protein_coding |  |  |  |  |  |  |  |
| TP53I3 | ENSG00000115129 | 2 | 24300303 | 24308731 | protein_coding |  |  |  |  |  |  |  |
| TP53TG1 | ENSG00000182165 | 7 | 86954541 | 86974831 | lincRNA |  |  |  |  |  |  |  |
| TRAFD1 | ENSG00000135148 | 12 | 112563305 | 112591407 | protein_coding |  |  |  |  |  |  |  |
| TRIM44 | ENSG00000166326 | 11 | 35684353 | 35829775 | protein_coding |  |  |  |  |  |  |  |
| TRIP12 | ENSG00000153827 | 2 | 230628554 | 230787955 | protein_coding | ✓ |  |  | ✓ |  |  |  |
| TRPC5 | ENSG00000072315 | X | 111017543 | 111326004 | protein_coding |  |  |  |  |  |  |  |
| TTBK2 | ENSG00000128881 | 15 | 43030932 | 43213007 | protein_coding |  |  |  |  |  |  |  |
| TTC3P1 | ENSG00000215105 | X | 74960541 | 74966749 | pseudogene |  |  |  |  |  |  |  |
| TTPAL | ENSG00000124120 | 20 | 43104526 | 43123244 | protein_coding |  |  |  |  |  |  |  |
| UBE2H | ENSG00000186591 | 7 | 129470572 | 129592789 | protein_coding | ✓ |  | ✓ | ✓ |  |  |  |
| UNC80 | ENSG00000144406 | 2 | 210636717 | 210864024 | protein_coding |  |  |  |  |  | ✓ |  |
| USP13 | ENSG00000058056 | 3 | 179370543 | 179507189 | protein_coding |  |  |  |  |  |  |  |
| USP32 | ENSG00000170832 | 17 | 58256455 | 58499831 | protein_coding |  |  |  |  |  |  |  |
| USP7 | ENSG00000187555 | 16 | 8985951 | 9058371 | protein_coding |  |  |  |  |  |  |  |
| USP9X | ENSG00000124486 | X | 40944888 | 41095832 | protein_coding |  |  |  |  |  |  |  |
| VAPB | ENSG00000124164 | 20 | 56964178 | 57026157 | protein_coding |  |  |  |  |  |  |  |
| WDR45B | ENSG00000141580 | 17 | 80572438 | 80606429 | protein_coding |  |  |  |  |  |  |  |
| ZNF274 | ENSG00000171606 | 19 | 58694396 | 58724928 | protein_coding |  |  |  |  |  |  |  |
| ZNF688 | ENSG00000229809 | 16 | 30580667 | 30584055 | protein_coding |  |  |  |  |  |  |  |
|  | ENSG00000232825 | 1 | 99469832 | 99614408 | antisense |  |  |  |  |  |  |  |
|  | ENSG00000260804 | 2 | 217081768 | 217084915 | antisense |  |  |  |  |  |  |  |
|  | ENSG00000203280 | 22 | 25498407 | 25508659 | lincRNA |  |  |  |  |  |  |  |

**Table S5. Experimental validation design.** Relative quantities were obtained by normalization on U6 RNA. Each condition is in biological triplicate, each of which is in technical duplicate for qPCR. Values are means± SD n=3 biological replicates by condition.

|  | mmu-miR-137-3p RQ geo mean |
| --- | --- |
| **1st batch** | |
| Control (ctrl, PX459 V2.0 empty vector) | 0.011±0.002 |
| Strong overexpression (OE .75) | 0.54±0.099 |
| G3 Knock out (KO) | 0±0 |
| **2nd batch** | |
| Control (ctrl, PcDNA3.2 empty vector) | 0.008±0.001 |
| Mild overexpression (OE .15) | 0.15±0.02 |
| Strong overexpression (OE .75) | 0.571±0.044 |

**Table S6. SNP weights.** The first column reports the SNP indicator; the second, the number of alleles other than the major homozygous (always represented as genotype “0”); the third reports the weight measured as A’ of the genotypic population considered relative to the major homozygous population.

| **snp** | **genotype** | **weight** |
| --- | --- | --- |
| rs12900014 | 0 | 0.500 |
| rs12900014 | 1 | 0.585 |
| rs12900014 | 2 | 0.875 |
| rs11033300 | 0 | 0.500 |
| rs11033300 | 1 | 0.683 |
| rs4941389 | 0 | 0.500 |
| rs4941389 | 1 | 0.368 |
| rs4941389 | 2 | 0.266 |
| rs11125946 | 0 | 0.500 |
| rs11125946 | 1 | 0.327 |
| rs11125946 | 2 | 0.285 |
| rs2853047 | 0 | 0.500 |
| rs2853047 | 1 | 0.305 |
| rs521877 | 0 | 0.500 |
| rs521877 | 1 | 0.361 |
| rs521877 | 2 | 0.242 |
| rs10012741 | 0 | 0.500 |
| rs10012741 | 1 | 0.665 |
| rs12582273 | 0 | 0.500 |
| rs12582273 | 1 | 0.651 |
| rs12582273 | 2 | 0.713 |
| rs8100664 | 0 | 0.500 |
| rs8100664 | 1 | 0.310 |
| rs8100664 | 2 | 0.297 |
| rs5008124 | 0 | 0.500 |
| rs5008124 | 1 | 0.642 |
| rs5008124 | 2 | 0.747 |
| rs4378203 | 0 | 0.500 |
| rs4378203 | 1 | 0.624 |
| rs4378203 | 2 | 0.706 |
| rs9544434 | 0 | 0.500 |
| rs9544434 | 1 | 0.386 |
| rs9544434 | 2 | 0.298 |
| rs334548 | 0 | 0.500 |
| rs334548 | 1 | 0.615 |
| rs334548 | 2 | 0.721 |
| rs3912022 | 0 | 0.500 |
| rs3912022 | 1 | 0.626 |
| rs3912022 | 2 | 0.741 |
| rs7946758 | 0 | 0.500 |
| rs7946758 | 1 | 0.627 |
| rs7946758 | 2 | 0.701 |

**Table S7. fMRI statistics.** Abbreviations: Amy: Amygdala; BA= Brodmann area; coor= coordinates in MNI space; k= cluster extent; L/R= Left or Right hemisphere; P = peak voxel whole-brain FWE-corrected p value (SVC in the replication sample).

| **PCI_miR-137_ – EP activity (discovery sample)** | | | | |  |  |  |  |  |
| --- | --- | --- | --- | --- | --- | --- | --- | --- | --- |
| **X coor** | **Y coor** | **Z coor** | **L/R** | **Gyrus** | **BA** | **k** | **t** | **Z** | ***P*** |
| 50 | 12 | 42 | R | Inferior Frontal Gyrus | 9 | 97 | 5.3 | 5.1 | 0.002 |
| **PCI_miR-137_ – EP activity (replication sample)** | | | | | | | | | |
| **X coor** | **Y coor** | **Z coor** | **L/R** | **Gyrus** | **BA** | **k** | **t** | **Z** | ***P*** |
| 46 | 12 | 31 | R | Inferior Frontal Gyrus | 9 | 42 | 3.6 | 3.5 | 0.007 |
| **PRS_miR-137_ – WM activity (discovery sample)** | | | | | | | | | |
| **X coor** | **Y coor** | **Z coor** | **L/R** | **Gyrus** | **BA** | **k** | **t** | **Z** | ***P*** |
| 27 | 27 | 58 | R | Superior Frontal Gyrus | 8 | 37 | 4.4 | 4.3 | 0.038 |
| **Complementary PRSscz - PRSmiR_-137_ – EP activity (discovery sample)** | | | | | | | | | |
| **X coor** | **Y coor** | **Z coor** | **L/R** | **Gyrus** | **BA** | **k** | **t** | **Z** | ***P*** |
| 27 | 8 | -29 | R | Parahippocampal/Amy | - | 16 | 4.2 | 4.1 | 0.05 |
| **Complementary PRSmiR_-137_ *no Darkorange* –WM activity (discovery sample)** | | | | | | | | | |
| **X coor** | **Y coor** | **Z coor** | **L/R** | **Gyrus** | **BA** | **k** | **t** | **Z** | ***P*** |
| 27 | 27 | 58 | R | Superior Frontal Gyrus | 8 | 30 | 4.4 | 4.5 | 0.01 |
